# TopoMetry systematically learns and evaluates the latent geometry of single-cell data

**DOI:** 10.1101/2022.03.14.484134

**Authors:** David S. Oliveira, Ana I. Domingos, Licio A. Velloso

**Affiliations:** University of Oxford, UK; University of Campinas, Brazil

**Keywords:** single-cell genomics, dimensional reduction, manifold learning, clustering, visualization

## Abstract

Reconstructing and investigating the geometry underlying data is a fundamental task in single-cell analysis, yet no unified framework exists for learning, evaluating, and diagnosing representations that faithfully preserve it. We present **TopoMetry**, a geometry-aware framework that learns intrinsic coordinate systems directly from the data and refines them into high-fidelity *spectral scaffolds*. These scaffolds capture both local neighborhoods and global structure, supporting downstream analysis such as clustering and visualization. In benchmarks across diverse single-cell datasets, TopoMetry preserved geometry more reliably than standard workflows and revealed biological signals otherwise obscured, including unexpected transcriptional diversity among T cells and links between RNA-defined subpopulations and clonal expansion. The full analysis can be executed with a single line of code to generate a comprehensive report, making the framework both powerful and accessible. Beyond individual findings, TopoMetry warrants a shift of focus from static two-dimensional projections to the systematic learning and evaluation of geometry itself, enabling more accurate exploration of cellular diversity.

## 1 Introduction

Single-cell genomics has enabled the systematic profiling of thousands to millions of individual cells, providing unprecedented insight into the diversity of cell types and states across tissues and organisms. These technologies generate high-dimensional data in which each cell is represented by measurements across tens of thousands of features, such as gene expression in single-cell RNA-seq (scRNAseq). Interpreting such data depends critically on computational analysis: the algorithms used to represent and compare cells directly shape the clusters, trajectories, and biological hypotheses that emerge from single-cell studies.

The prevailing analytical workflow begins by reducing dimensionality with Principal Component Analysis (PCA)^1^, followed by the construction of a neighborhood graph for Leiden clustering and visualization with algorithms such as UMAP (Uniform Manifold Approximation and Projection)^2,3^. The “PCA → neighborhood graph → Leiden clustering and UMAP” pipeline has become the *de facto* standard in single-cell genomics^3–5^ as it was adopted by popular toolkits such as `SCANPY` and `Seurat` and underpins a wide range of downstream tasks, from cell annotation to trajectory inference and dataset integration. Its popularity reflects both its computational efficiency and its capacity to produce intuitive visual summaries of cellular heterogeneity.

Despite its wide adoption, the PCA-based workflow rests on assumptions that are difficult to verify for most single-cell datasets^1,2^. The use of PCA presumes that cell states can be represented as linear combinations of genes and that biological variation is captured by global variance (an assumption also shared by variational models such as scVI; Supplementary Figure S1a). UMAP assumes that cells are *uniformly* sampled from a manifold with a *constant* local metric. Violations of these assumptions lead to similarity graphs and embeddings that distort cellular relationships, with direct consequences for clustering outcomes, lineage reconstructions, and biological interpretation of results. In addition, any two-dimensional projection will inevitably introduce distortions, which has fueled debate about how much trust should be placed in embeddings obtained with tools such as UMAP^6^.

The field currently lacks a systematic evaluation of the standard PCA-based pipeline and its impact on the geometric fidelity of single-cell analyses. While benchmarking studies have compared clustering or integration algorithms^7,8^, none have assessed whether the obtained representations preserve the original manifold structure that encodes cellular identities. In particular, there has been no unified framework for quantifying how neighborhood graphs, diffusion processes, and embeddings diverge from the ground-truth geometry. Without such evaluation, distortions introduced at early stages of the pipeline may propagate through analyses, affecting biological conclusions in ways that are difficult to detect or correct.

Ideally, the field should adopt a theoretical framework for modeling single-cell data that minimizes assumptions while providing stable and intuitive representations across diverse datasets. Such a framework should (i) treat single-cell data as samples drawn from manifolds with heterogeneous and potentially disconnected supports, (ii) explicitly estimate intrinsic dimensionalities rather than fixing representation size arbitrarily, and (iii) evaluate how well similarity graphs and embeddings preserve the multiscale geometry of the data. By grounding analyses in rigorous geometric principles, such a framework would enable more reliable inference of cell states, lineages, and transitions (Supplementary Figure S1b).

To address this gap, we developed TopoMetry, a framework for single-cell geometric analysis designed to preserve intrinsic data structure independently of distributional or sampling artifacts. TopoMetry considers the existence of multiple manifolds in each single-cell dataset, each representing a distinct macro-population or lineage, and decomposes the joint geometry of the data into a *spectral scaffold*: a set of hundreds of components that, analogous to the harmonics of a Fourier transform, collectively capture the global and local structure of the dataset. A refined cell–cell similarity graph is then constructed from this scaffold and used for downstream analyses. TopoMetry also introduces new geometry preservation metrics and distortion visualization tools, making it possible to quantitatively and qualitatively evaluate latent representations and 2-D visualizations.

We evaluated TopoMetry against the prevailing PCA-based standard (Supplementary Figure S1c) in an extensive collection of single-cell datasets. Across all datasets, the proposed workflow yielded representations that better capture the original data geometry and clustering results that are better suited for the detection of rare cell populations. We found that PCA’s poor performance in preserving geometry was associated with its failure to explain variance, highlighting its inappropriateness for single-cell analyses for the first time. The difference in performance between the PCA-based and TopoMetry workflows was particularly noticeable in datasets containing T cells, in which TopoMetry identifies a large number of T cell clusters that are completely missed by the standard workflow. Using paired TCR and RNA data, we explored the geometrical properties of clonal expansion in single-cell data and found that these additional clusters were associated with TCR clonality. Collectively, our results demonstrate the superiority of a geometric framework for single-cell analysis over the prevailing standards.

## 2 Results

### 2.1 A framework to systematically learn and evaluate manifolds in single-cell data

To study cellular diversity, single-cell data are often represented as points scattered across a high-dimensional space of measured features (e.g., gene expression in scRNAseq). The challenge is to uncover the hidden structure — the manifold of cell identities — that explains how these points are organized into cell types, lineages, and states. Our goal with TopoMetry is to provide a framework that learns this manifold directly from the data and builds on its geometrical properties, without relying on restrictive assumptions on its topology, distribution, or intrinsic dimensionality.

TopoMetry takes as input a scaled (standardized or Z-score normalized) matrix of features per cell (e.g., genes per cell). It then builds a neighborhood graph that connects each cell to its *k* most similar counterparts, which is transformed into a similarity matrix using decay-adaptive, manifold-aware kernels that account for local intrinsic dimensionality and sampling density (Figure 1a). These kernels reduce density-driven bias and support the construction of Laplacian-type and diffusion operators that approximate manifold geometry^9–12^ (Figure 1a, Supplementary Figure S1d)).

**Figure 1:**
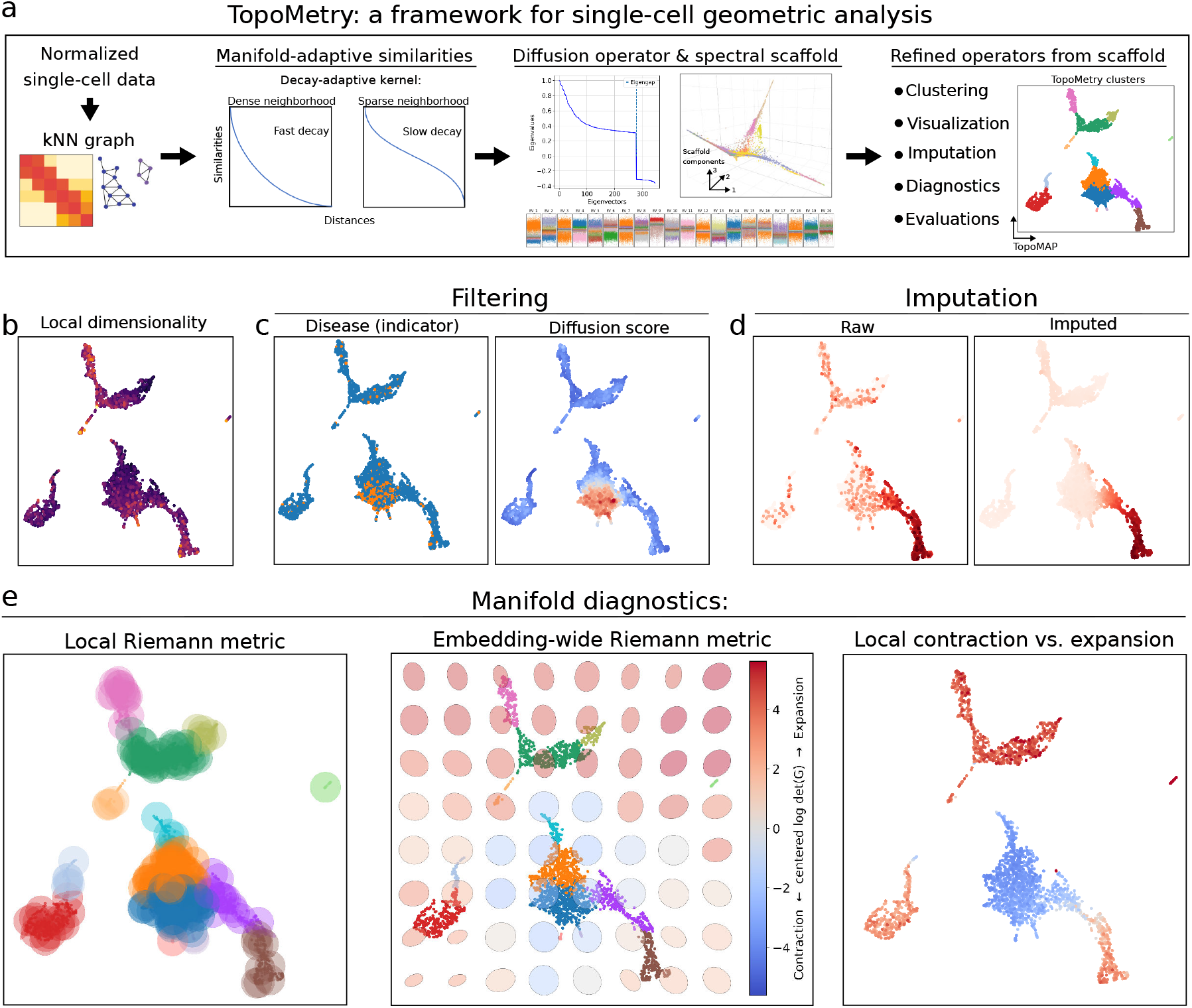
A framework for single-cell geometric analysis. (a) Schematic overview of the TopoMetry algorithm. From an input single-cell dataset (e.g., normalized and scaled scRNAseq data), TopoMetry builds a kNN graph, which is used to learn manifold-adaptive similarities with a decay-adaptive kernel suitable for constructing Laplacian-type and diffusion operators. After estimation of intrinsic dimensionality, these operators are decomposed into a spectral scaffold with up to hundreds of components that jointly explain all of the underlying geometry of the dataset. The spectral scaffolds are used to learn refined Laplacian-type and diffusion operators of the scaffolds themselves, encoding “the geometry of the geometry”. The scaffolds and operators constitute key TopoMetry outputs and can be utilized for downstream tasks, such as clustering, visualization, imputation, evaluation, and diagnostics, in a geometry-aware manner. TopoMetry utilities include (b) estimation of local intrinsic dimensionality, (c) filtering of categorical signals, and (d) imputation and denoising. Crucially, TopoMetry introduces the visualization of manifold diagnostics (e) for single-cell data, in which distortions induced by 2-D embeddings can be identified and investigated from a local, global, and contraction/expansion perspective.

TopoMetry proceeds by eigendecomposing these operators into hundreds of orthogonal components that together define a *spectral scaffold* (Figure 1a). Each component serves as a latent coordinate that captures a distinct mode of variation. A *multiscale spectral scaffold* aggregates coordinates across diffusion times to reconcile local neighborhoods and long-range global organization. The number of scaffold components is determined automatically by estimating intrinsic dimensionality (I.D.), rather than fixed *a priori*^13,14^ (Figure 1b). The scaffold thus constitutes a latent space that encodes the geometrical properties of the data, is robust to different neighborhood sizes (Suppl. Figure S1e), and supports the construction of refined similarity graphs and diffusion operators from the scaffold itself, thus capturing the “geometry of the geometry”. These refined graphs and operators serve as high-fidelity inputs for downstream tasks such as clustering and visualization (Figure 1a), and can also be exploited to evaluate how information propagates along the manifold, enabling graph-based filtering of categorical or continuous signals (Figure 1c) and data-smoothing operations such as imputation and denoising (Figure 1d). Finally, TopoMetry leverages the Riemannian metric to provide manifold diagnostics that reveal contraction, expansion, and local distortions introduced by two-dimensional maps, allowing geometry-aware interpretation (Figure 1e)^15^.

The proposed approach has several conceptual advantages over current standards. First, it assumes only that samples lie approximately on manifolds—a minimalistic and yet biologically meaningful premise consistent with the Waddington epigenetic landscape^16^ and already implicit in many downstream analyses such as lineage inference and pseudotime estimation. Second, it eliminates the need to pre-select the number of components by estimating I.D. directly and identifying gaps in the scaffold eigenspectrum (Figure 1a). Third, it evaluates fidelity with quantitative metrics rather than relying solely on qualitative visualizations. Finally, by coupling manifold-based analysis with modern graph-layout techniques (e.g., UMAP^2^, PaCMAP^17^), TopoMetry moves beyond fixed pipelines toward systematic, data-driven discovery. The full analysis can be executed with a single line of code, generating a comprehensive report while remaining fully customizable and compatible with `AnnData/Scanpy`^4^ and the broader *Python* single-cell ecosystem. In summary, TopoMetry brings manifold learning and geometric analysis into single-cell genomics in a way that is rigorous, computationally efficient (Supplementary Figure S1f), and accessible to researchers from all backgrounds.

### 2.2 Evaluating geometry-preservation across representations

A central question in single-cell analysis is whether a low-dimensional representation preserves the original geometry of cellular relationships, such as multiple cellular lineages cohabiting the same gene expression space. Such geometry is encoded by the diffusion operator built from the initial neighborhood graph^10,18^. To evaluate geometry preservation, we introduce a family of *operator-native* metrics that compare representations at the level of their learned diffusion kernels/transition operators rather than at the level of raw Euclidean coordinates. This choice is deliberate: many downstream tasks (visualization, clustering, trajectories, pseudotime) implicitly act on a graph or Markov operator, so fidelity should be judged in that native space.

First, we quantify local neighborhood agreement in two complementary ways. The **Sparse Neighborhood F1** (P-F1@k) measures set overlap between the top-*k* transition supports of two operators (insensitive to weights, sensitive to who the neighbors are). The **row-wise Jensen–Shannon similarity** (`P-JS`) compares each row as a probability distribution over neighbors (sensitive to weights, i.e., how strongly neighbors are connected). Together, P-F1@k and `P-JS` capture whether local neighborhoods are preserved *and* whether their transition probabilities remain consistent. Second, we assess meso-to global-scale geometry using diffusion coordinates. The **Spectral Procrustes** score (`SP`) aligns truncated diffusion maps (at multiple diffusion times) via orthogonal Procrustes and reports an *R*^2^ goodness-of-fit; high values indicate that two operators induce essentially the same geometry up to a rotation. Collectively, these measures probe complementary facets of fidelity: neighborhood composition, edge weights, global alignment, and connectivity structure. Importantly, all metrics compare the diffusion operator of the projection built by each method to the native diffusion operator of normalized gene expression, ensuring a fair, task-relevant comparison independent of a particular 2-D layout.

#### Benchmark datasets

To apply the defined geometry preservation metrics to representations learned with the current standard workflow or TopoMetry, we curated a collection of 68 scRNAseq datasets spanning multiple organs, tissues, and species (humans and mice). The corpus comprises atlas-scale datasets with a complex hierarchical structure, as well as focused studies on rare or transitional populations, thereby representing single-cell genomics data in general (Figure 2a, Suppl. Table 1).

**Figure 2:**
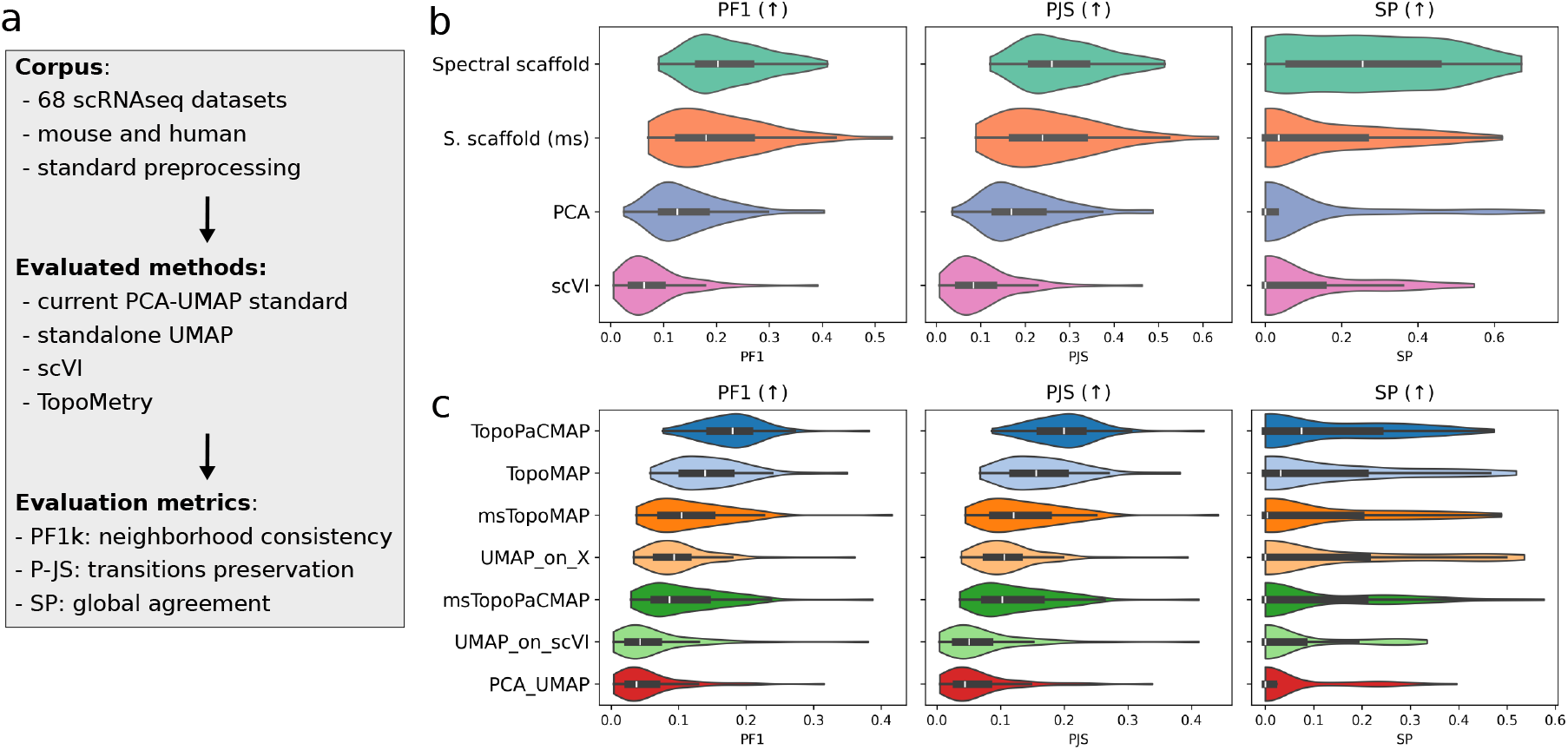
Geometry preservation benchmark. (a) Schematic representation of the benchmark workflow, in which a corpus composed of 68 scRNAseq datasets was collected, preprocessed, and analyzed with i) the current PCA→UMAP standard, ii) standalone UMAP (graph from high-dimensional gene expression space), iii) scVI (a popular tool for variational inference), and iv) TopoMetry. (b) Violin plots representing geometry-preservation metrics for lower-dimensional latent spaces learned with PCA and scVI, compared to TopoMetry’s spectral scaffolds. TopoMetry’s scaffolds achieved systematically higher scores across all metrics. (c) Violin plots representing geometry-preservation metrics for 2-D visualizations obtained with the evaluated methods. Except for PaCMAP on TopoMetry’s multiscale spectral scaffold, the geometry-aware visualizations achieved systematically higher scores. Visualizations based on scVI and PCA latent space presented the lowest scores.

#### Evaluating workflows

We applied the metrics to representations produced by (i) the standard workflow (kNN graph built on the PCA space^1,19^), (ii) a “pure” standalone UMAP approach (kNN graph built directly from the high-dimensional space^2^), (iii) scVI (a widely used variational framework^20,21^), and (iv) TopoMetry (Figure 2a). We then computed the geometry preservation scores. For all analyzed datasets, the spectral scaffolds learned by TopoMetry scored consistently higher than the latent spaces learned by PCA or scVI (Figure 2b). In agreement with these findings, 2-D visualizations obtained with TopoMetry also performed better than those obtained with PCA or scVI (Figure 2c). Noticeably, “pure” UMAP visualizations had intermediary scores between TopoMetry and PCA→UMAPs, highlighting that PCA usage is not universally appropriate in single-cell genomics. In sum, these results indicate that TopoMetry’s manifold-centered approach yields representations that better preserve the original geometrical properties of single-cell data, thus being better suited for downstream analyses and biological interpretation.

Next, we further investigated the reasons underlying the poor performance of PCA-based graphs and visualizations. We found that PCA performance correlates with its ability to explain most of the original variance (Suppl. Fig. S2a). Surprisingly, the total variance explained by PCA was remarkably low across datasets (as little as 20%). Total explained variance decreased as the number of highly-variable genes (i.e., dimensionality) increased, and averaged only ∼36% across all datasets with default gene selection (Suppl. Fig. S2b). Failure to explain variance could not be attributed to insufficient components, as demonstrated by eigenspectra (Suppl. Fig. S2c) and cumulative explained variance curves (Suppl. Fig. S2d), where an *ad hoc* “elbow point” was consistently found around 30–50 components. Total explained variance was weakly associated with cell number (Suppl. Fig. S2e). Such low values of explained variance are widely known as a hallmark of highly non-linear systems in the broader machine-learning community^22,23^, albeit apparently unnoticed within the single-cell niche. Collectively, these observations suggest that PCA fails on single-cell data due to intrinsic properties, such as nonlinearity, which can be highly dataset-specific. These limitations highlight the need for approaches that move beyond variance-based assumptions and instead preserve manifold geometry directly.

### 2.3 TopoMetry resolves cellular development lineages in tiny and atlas-scale systems

A natural application of such geometry-aware representations is the reconstruction of developmental trajectories. To test this, we first applied TopoMetry to scRNAseq data from the developing murine pancreas, a well-established benchmark in RNA velocity studies^24,25^. Standard PCA→UMAP embeddings (Figure 3b, Suppl. Fig. S3a–b) identified broad lineages but failed to represent the cell cycle structure, placing mitotic cells into ambiguous positions along differentiation axes. In contrast, TopoMetry projections (TopoMAP, Figure 3a,c) reconstructed a closed-loop geometry of the cell cycle and positioned proliferating cells alongside other cycling populations, faithfully reflecting the underlying manifold. RNA velocity streamlines confirmed the accuracy of this representation (Figure 3d), while the PCA-based embedding produced vector fields inconsistent with known lineage relationships (e.g. epsilon cells giving rise to beta cells^26^). Overlaying individual scaffold components on TopoMAP projections further showed that each component captures a distinct facet of the manifold, from global developmental trajectories to localized cellular states (Suppl. Fig. S3c).

**Figure 3:**
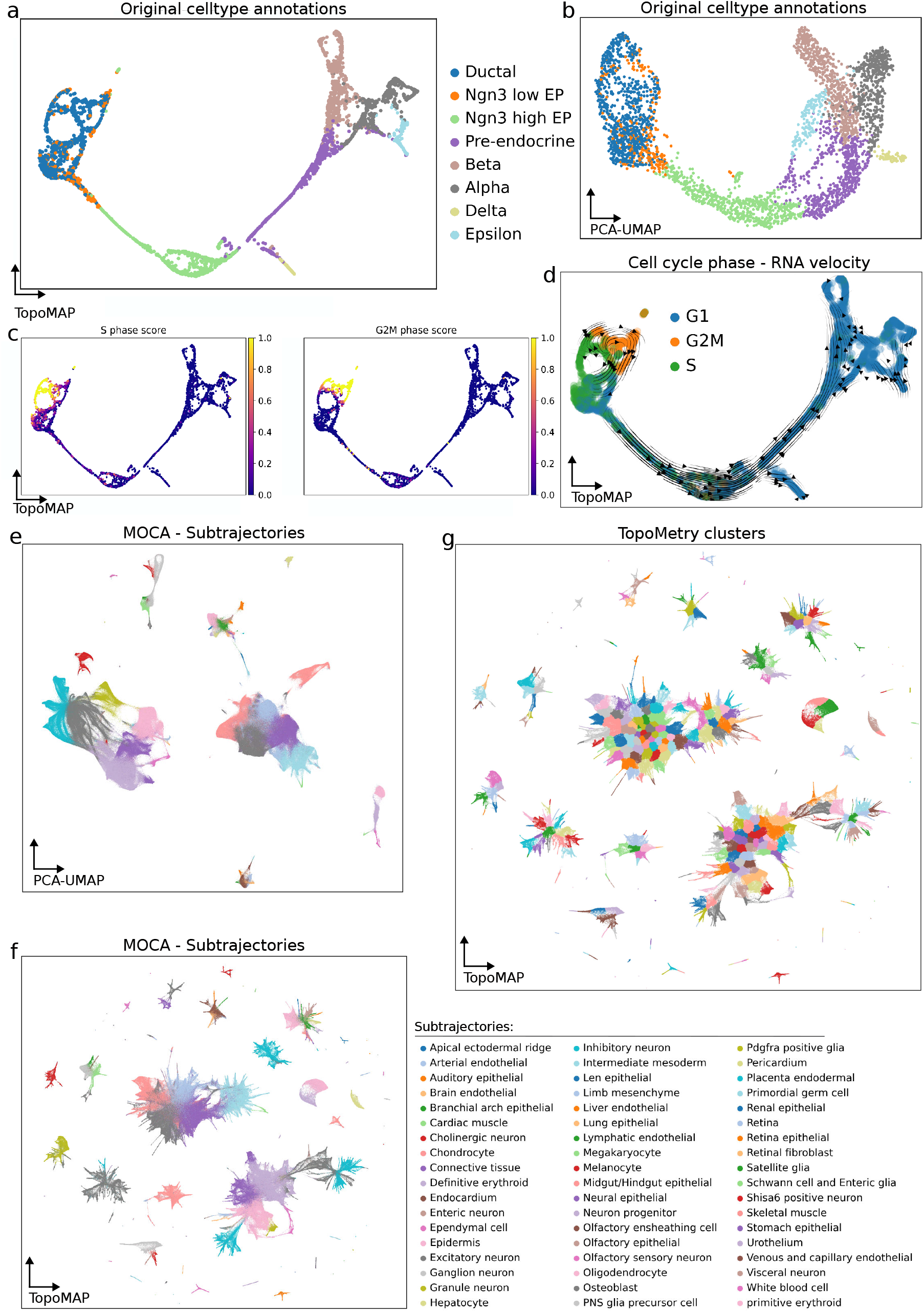
Inferring cellular lineages with TopoMetry. (a) TopoMAP and (b) PCA→UMAP visualizations of the Pancreas dataset showing cellular developmental trajectories in the murine pancreas, colored by original cell type annotations. (c) TopoMAP visualization, colored by inferred scores of different phases of the cell cycle, and (d) the predicted phase for each cell with RNA velocity overlay. Note how RNA velocity trajectories largely agree with the identified cell cycle structure and the represented geometry. (e) PCA→UMAP visualization of the Mouse Organogenesis Cell Atlas (MOCA), comprising ∼1.3 million cells collected during murine embryo development, colored by refined subtrajectories annotation. (f–g) TopoMAP visualizations of MOCA, colored by original annotations on refined subtrajectories (f), and TopoMetry’s clustering results (g). Note how the TopoMAP embedding successfully separates main and refined trajectories and adds enhanced detail and resolution on the diversity of subpopulations arising during development.

We demonstrated that TopoMetry can scale to organism-wide atlases by analyzing the Mouse Organogenesis Cell Atlas (MOCA), a compendium of ∼1.3 million cells spanning murine embryonic development^27^. Both PCA→UMAP and TopoMetry separated the originally annotated differentiation trajectories and cell types into distinct regions of the manifold (Figure 3e–f). However, TopoMetry revealed a much richer hierarchy of sub-trajectories, capturing ∼380 subpopulations compared to the 56 originally described (Figure 3g). These included a particularly fine-grained resolution of neuronal lineages, consistent with the known diversity of the developing nervous system. By encoding long-range and local geometry simultaneously, the TopoMAP embedding reconstructed refined developmental paths that remained unresolved in PCA-based visualizations.

Together, these analyses show that TopoMetry faithfully represents cellular dynamics across scales: from local cycles and branching lineages in small datasets to hundreds of coexisting trajectories in million-cell atlases. These capabilities make TopoMetry an effective framework for studying lineage inference in direct connection with the Waddington epigenetic landscape^16^.

### 2.4 TopoMetry unveils unexpected transcriptional diversity of T cells

When applying TopoMetry to public single-cell datasets, we repeatedly observed an unexpected pattern in peripheral blood mononuclear cell (PBMC) data: TopoMetry consistently revealed a much greater diversity of T cells than the standard PCA→UMAP workflow. Such pattern first became evident in the widely used *pbmc68k* dataset, which contains ∼68,000 PBMCs from a healthy donor. While TopoMetry uncovered an unexpectedly fine-grained landscape with close to one hundred distinct T cell clusters (Figure 4a), PCA-based UMAP represented T cells as a few broad clusters (Figure 4b). Importantly, major immune cell classes were similarly well separated in both approaches, underscoring that the difference lies specifically in the resolution of T cell diversity.

**Figure 4:**
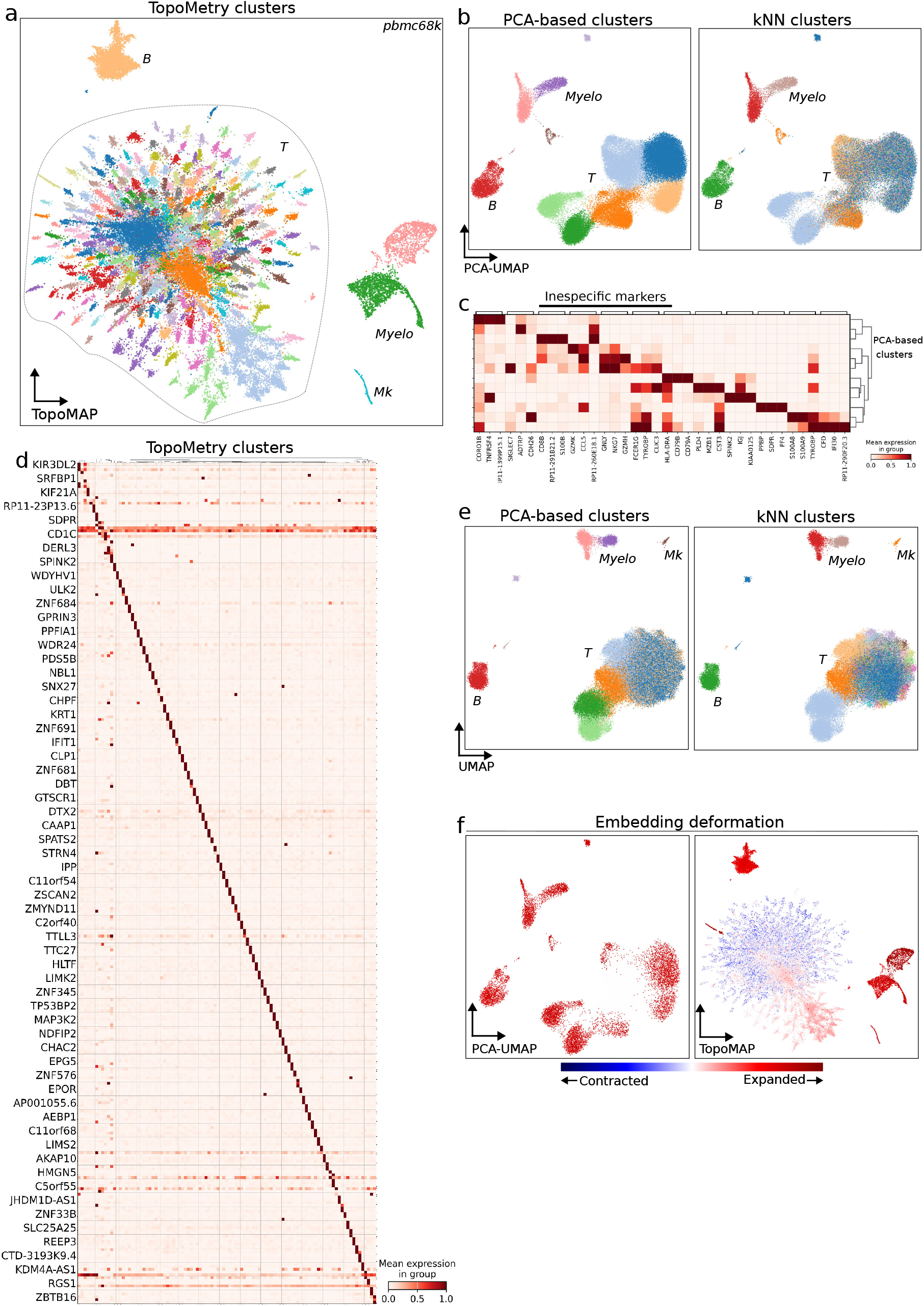
TopoMetry unveils unexpected transcriptional diversity of T cells. Analysis of the pbmc68k dataset, comprising approximately 68,000 peripheral blood mononuclear cells from a healthy donor (10X Genomics). (a) TopoMAP visualization colored by TopoMetry’s clustering results. Main cell types are well separated, and T cells present an unexpected high diversity, with approximately a hundred clusters identifying T cell subpopulations. (b) Standard PCA→UMAP visualizations colored by clustering results obtained with the PCA-derived graph (left) and the kNN graph from the high-dimensional gene expression space (right), presenting the same global separation of main cell types but disagreeing on T cells. (c) Matrixplot of the top 3 marker genes found for PCA-based clusters, highlighting the presence of non-specific markers for T cells. (d) Matrixplot of the top 3 marker genes found for TopoMetry clusters, presenting highly specific marker expression. (e) Standalone UMAP visualizations of the same data, colored by PCA-based (left) and kNN-based clustering results (right). Note how the standalone approach detects part, but not all, of the T cell clusters identified with TopoMetry. (f) Contraction/expansion diagnostics of PCA→UMAP (left) and TopoMAP (right) visualizations. Note how the PCA→UMAP approach expands most regions of the cell identity manifold, while TopoMAP contracts the region inhabited by T CD4 lymphocytes when projecting TopoMetry’s refined graphs to a 2-D space.

The discrepancy was further highlighted when examining marker gene expression. T cell clusters obtained with PCA-based graphs presented non-specific markers (Figure 4c), whereas TopoMetry clusters exhibited highly specific expression signatures (Figure 4d). These included canonical T cell markers, interleukin receptors, and T cell receptor (TCR) genes. Such specificity indicates that TopoMetry is capturing biologically meaningful substructure rather than technical noise. Additional TopoMetry projections consistently separated these substructures (Suppl. Fig. S4a).

Independent workflows corroborated these findings. A “pure” UMAP approach, in which graph construction is performed directly on the high-dimensional gene expression matrix, recovered a subset of the additional T cell substructures but not their full extent (Figure 4e). Similarly, clustering on a standalone kNN graph without PCA partially agreed with TopoMetry’s results (Figure 4e, Suppl. Fig. S4b). These comparisons suggest that the additional diversity is present in the original data but systematically missed when PCA is imposed as the first step of the analysis.

We next asked whether technical artifacts could explain these unexpected T cell clusters. Doublet detection using Scrublet^28^ showed no doublet enrichment in these clusters (Suppl. Fig. S4c–d), arguing against artifactual origins. Finally, distortion diagnostics revealed that the PCA→UMAP embedding expands large portions of the T cell manifold, while TopoMetry’s refined graphs contract the CD4^+^ T cell region (Figure 4f), consistent with genuine underlying structure rather than projection artifacts. Together, these results establish that TopoMetry reliably detects transcriptional heterogeneity in T cells that is largely obscured by conventional workflows.

The fine-grained T cell structure revealed in pbmc68k can be further dissected through the spectral scaffold itself. When visualizing the first 40 scaffold components, the earliest components captured global immune class separations and long-range relationships, while subsequent components progressively refined local features and delineated specific T cell subsets (Suppl. Fig. S5). This decomposition illustrates how TopoMetry resolves the manifold region by region, with each component encoding a distinct aspect of cellular heterogeneity.

We next examined whether these unexpected T cell clusters were also found in PBMCs sampled from disease contexts. In case studies of systemic lupus erythematosus, dengue fever, and multiple sclerosis, the standard PCA→UMAP workflow produced only broad T cell clusters with largely non-specific markers (Suppl. Fig. S6a–c). Standalone UMAP and kNN clustering recovered a subset of these additional populations, but TopoMetry consistently revealed the full breadth of T cell diversity, detecting numerous clusters supported by highly specific marker gene expression. These results suggest that the richer representation of T cells is not dataset-specific but extends across diverse biological contexts.

A closer inspection of marker signatures indicated that the identities of these T cell clusters varied across donors and conditions. Some clusters reflected canonical CD4 or CD8 states, whereas activation signatures or effector molecules distinguished others. This variability points to genuine donor-specific biology rather than technical artifacts. In particular, the heterogeneity is consistent with the highly individualized T cell receptor (TCR) repertoire: each individual carries a unique set of clonotypes, which could lead to subtle transcriptional imprints that are easily masked by linear or density-biased workflows. Additionally, recent studies suggest that T cells sharing the same TCR clonotype (or recognizing the same epitope) tend to display similar transcriptional phenotypes, and joint representations of both RNA and TCR data more clearly distinguish antigen-specific subpopulations than those based only on the transcriptome^29–31^. These observations motivated us to test whether the additional T cell resolution provided by TopoMetry is associated with TCR clonotypes.

### 2.5 TopoMetry resolves clonal dynamics of T cells from RNA expression

To investigate if the additional clusters of T cells exclusively discovered by TopoMetry are associated with TCR clonal dynamics, we analyzed two public datasets where scRNAseq was paired with TCR sequencing: the ECCITE-TCR study of CD8^+^ T cell responses to SARS-CoV-2 vaccination and infection^30^, and the T cell compartment of the Tissue Immune Cell Atlas (TICA)^32^, which combines scRNAseq and VDJ-seq across multiple human tissues.

#### SARS-CoV-2 vaccination data

In the ECCITE-TCR dataset, TopoMetry once again revealed a rich diversity of CD8^+^ subpopulations (Fig. 5a) that were absent from PCA→UMAP representations (Suppl. Fig. S7a) and presented highly specific markers (Fig. 5b). Many of these additional clusters corresponded to effector memory (*Tem*) and central memory (*Tcm*) populations, as indicated by original cell type annotations (Fig. 5c). These clusters were associated with less frequent clonotypes (Fig. 5d), suggesting that the transcriptional variation uncovered by TopoMetry reflects underlying clonal dynamics. Cell-cycle trajectories of proliferating antigen-specific CD8^+^ lymphocytes were also faithfully encoded by TopoMetry representations, producing coherent loop-like structures in 2-D projections (Fig. 5e), whereas the same geometry was broken or distorted in PCA-based results even with the use of joint RNA–TCR representations (Suppl. Fig. S7a–g). Beyond visualizations, the spectral scaffold learned by TopoMetry captured both global lineage structure and local expansions (Suppl. Fig. S8a), while estimates of local intrinsic dimensionality remained stable across the manifold (Suppl. Fig. S8b). Probability density mapping of clone sizes on the TopoMetry embedding further confirmed that small and rare clonotypes localized to the additional TCM/TEM clusters, while hyperexpanded clones occupied distinct regions of the manifold (Suppl. Fig. S8c).

**Figure 5:**
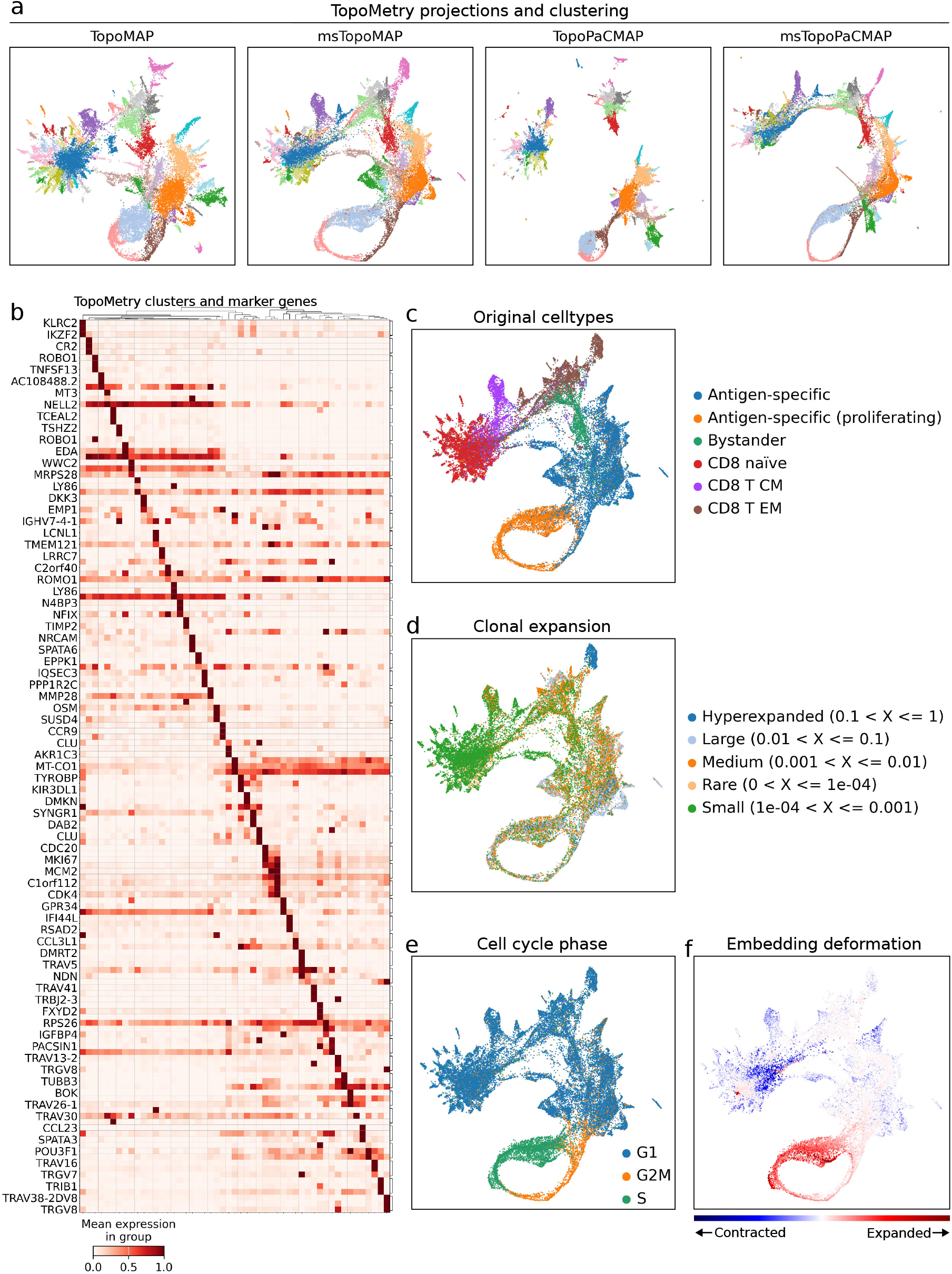
TopoMetry detects T cell clonal expansion dynamics from gene expression. Analysis of the ECCITE-TCR dataset, comprising circulating T CD8^+^ lymphocytes collected from human donors in baseline conditions and after SARS-CoV-2 vaccination or infection. (a) TopoMetry’s default visualizations, colored by TopoMetry’s clustering results. Note how projections derived from the fixed-time scaffold better preserve the local structure of the dataset, while projections derived from the multiscale scaffold better preserve long-range relationships and overall global structure. Despite minor differences, all visualizations correctly represent the cell cycle geometry of proliferating lymphocytes and the transcriptional diversity of central (TCM) and effector memory (TEM) lymphocytes. (b) Matrixplot of the top 3 marker genes found for TopoMetry clusters, presenting highly specific marker expression. (c–f) TopoMAP visualizations colored by original cell type annotations (c), clonal expansion information (d), predicted phase of the cell cycle (e), and contraction/expansion diagnostics. Note how the small clusters of TCM and TEM correspond to smaller clonotypes (ranging from small to medium), how the identified cell cycle geometry agrees with cell cycle predictions, and how the former are contracted while the latter are expanded in the 2-D visualization.

#### Tissue Immune Cell Atlas

We next examined the TICA dataset^32^, comprising paired RNA and VDJ profiles of human T cells across tissues and donors. As in previous analyses, TopoMetry identified numerous clusters within the CD4^+^ compartment that were either merged or absent in both the PCA-based workflow and original cell type annotations (Fig. S9a-b). Using the `scirpy` TCR analysis toolkit (^33^), we queried epitope databases for antigens matching the TCR repertoire of this dataset, and found that the additional clusters uncovered by TopoMetry recognized antigens from specific species (Fig. S9c). Of note, we found that one of TopoMetry’s clusters corresponded to a single clonotype of tissue-resident memory T CD8^+^ cells with specificity for SARS-CoV-2 antigens (Fig. S9c). To further investigate clonal expansion dynamics, we visualized the 30 largest clonotypes with TopoMetry, and found a strong correspondence between these clonotypes and the fine-grained cluster structure identified by TopoMetry (Fig. S9c). Several of these clusters corresponded to hyperexpanded clones, including the population recognizing SARS-CoV-2 antigens (Fig. S9e). Clonal expansion analysis further confirmed that hyperexpanded clones mapped to distinct CD8^+^ and NK populations, whereas the TopoMetry-specific CD4^+^ clusters corresponded to clonotypes with modest expansion (Fig. S9f). Finally, to evaluate the specificity of the cluster-clonotype associations between workflows, we performed TCR repertoire overlap analysis, revealing that the PCA-based workflow and original cell type annotations resulted in strong repertoire overlap (Suppl. Fig. S9f). The “pure” standalone kNN workflow (Suppl. Fig. S9g) presented clearly weaker overlap, followed by TopoMetry (Suppl. Fig. S9h), which presented minimal overlap. These results demonstrate that TopoMetry’s clustering results achieve a strong association with clonotypes inferred from TCR sequence similarity, in contrast to the poor performance of the PCA-based approach and the intermediate performance of standalone kNN graphs.

Collectively, these results show that TopoMetry’s geometry-aware workflow uncovers transcriptional imprints of TCR clonotypes directly from RNA expression. Across both SARS-CoV-2 vaccination and atlas-scale datasets, the additional T cell clusters revealed by TopoMetry corresponded to clonally distinct populations, many of which were invisible to PCA-based workflows. This finding closes the loop on our incidental observation in PBMC data: the unexpected T cell diversity resolved by TopoMetry reflects the clonal architecture of the immune repertoire. More broadly, these analyses highlight how preserving geometry enables faithful recovery of biological structure, bridging transcriptional diversity with clonotype identity in a way that linear or density-biased methods fail to achieve.

## 3 Discussion

In this work, we introduced TopoMetry, a framework for geometry-aware analysis of single-cell data that systematically learns manifold structure directly from the data. By constructing similarity graphs with kernels adaptive to local density and intrinsic dimensionality, TopoMetry extracts *spectral scaffolds* that capture the full range of geometric variation present in cellular systems. These scaffolds serve as the basis for refined diffusion graphs that preserve both local neighborhoods and long-range relationships, enabling downstream tasks such as clustering, visualization, imputation, and lineage inference in a way that is faithful to the original geometry. Crucially, TopoMetry also provides operator-native fidelity scores and distortion diagnostics, allowing users to evaluate when and how embeddings diverge from the underlying manifold. Applications across diverse datasets demonstrated that TopoMetry yields stable, faithful, and interpretable representations, often revealing subtle yet biologically meaningful structures invisible to existing workflows. Most notably, TopoMetry uncovered an unexpectedly rich transcriptional landscape of T cells in peripheral blood, linked to TCR repertoires and clonal expansion, demonstrating the biological insights that become accessible when geometry is preserved.

The results presented here also expose fundamental flaws in the current PCA→UMAP standard. The reliance on PCA as a first analysis step imposes restrictive assumptions: that linear variance captures the relevant biological signal and that the dataset is globally well-approximated by a low-rank linear subspace^1,19^. Our systematic evaluation shows that these assumptions fail dramatically in single-cell genomics: PCA typically explains less than 40% of the variance in standard settings, and the variance explained decreases as the number of highly variable genes grows. These results mean that PCA discards the majority of biological signal before manifold learning even begins. The consequence is clear: PCA-based embeddings systematically collapse diverse cell populations into broad, poorly resolved clusters, as observed with datasets containing T cells, where lineage- and clonotype-associated subpopulations vanish into a single homogeneous cloud. UMAP then compounds the issue by optimizing the layout from the already distorted PCA-based neighborhood graph. The field has relied on this standard for years–not because it is faithful, but because it is convenient. Our results demonstrate that such convenience comes at the cost of accuracy, discarding most of the underlying information and obscuring biologically meaningful structure.

TopoMetry does not simply extend previous diffusion-based methods such as Diffusion Maps^10^ or PHATE^34^; it provides a systematic and general framework for geometric analysis and manifold learning in single-cell data. Diffusion Maps introduced the idea of capturing data geometry across scales, while PHATE emphasized visualization of diffusion potentials. TopoMetry unifies and generalizes these insights into a pipeline that is both rigorous and practical: adaptive kernels ensure robustness to uneven sampling, spectral scaffolds encode manifold structure at data-driven resolutions, refined operators capture “the geometry of the geometry,” geometry preservation metrics evaluate the learned representations, and Riemannian diagnostics offer principled tools for assessing distortion in visualizations. This combination of theory and practice positions TopoMetry as a distinct step forward. Importantly, its impact is already visible in the field: Hale *et al*. recently used TopoMetry to detect morphologically distinct T cell populations in a high-content imaging study published in *Science*^35^, and Tedeschi *et al*. integrated it into their analysis of multimodal single-cell atlases^36^. That independent groups are already leveraging TopoMetry underscores its utility and relevance.

As with any method, TopoMetry has limitations. Its current implementation is optimized for exploratory analysis rather than for online mapping of new cells, which restricts its use in scenarios requiring direct projection into precomputed manifolds. Computational cost is another consideration: although scalable to millions of cells, geometric analysis is still more computationally intensive than PCA. Finally, while our discovery of clonally linked T cell populations highlights the method’s ability to reveal new biology, these findings call for follow-up experimental validation. Encouragingly, studies employing joint representations learned from paired TCR and RNA data have identified populations that resemble those identified by TopoMetry^29,30^, indicating that these arise from reproducible biological signals.

Future work will extend TopoMetry’s geometric-centered vision in several directions. Integrating geometric autoencoders^37^ could enable inverse mapping and online embedding of new cells. Extensions to multimodal data are natural, particularly for integrating RNA, chromatin, protein, and receptor information into unified geometry-aware representations^38^. Biologically, TopoMetry opens new opportunities to study T cell clonality, lineage diversification, and other processes where subtle geometric variation encodes key biological signals. More broadly, the framework could be applied to any high-dimensional single-cell modality, from epigenomics to imaging cytometry, wherever faithful preservation of geometry matters.

In sum, TopoMetry represents a conceptual shift: a framework that places geometry at the center of single-cell analysis and a tool for discovery, capable of revealing structure even in datasets that have already been thoroughly examined by the community. The rich T cell diversity uncovered in PBMC data, long considered a “solved” dataset and ubiquitously used for tutorials and demonstrations, illustrates this power. Far from being exhausted, such datasets are valuable resources of potentially new biological insight if analyzed with appropriate geometric tools. TopoMetry provides these tools, and in doing so, it sets the stage for a new generation of geometry-aware single-cell genomics.

## 4 Methods

### 4.1 TopoMetry

### 4.2 TopoMetry: Topological geoMetry

#### Manifold hypothesis and sampling model

Let ℳ ⊂ ℝ^*D*^ be a compact, *d*-dimensional *C*^2^ Riemannian submanifold (possibly with boundary) with metric *g* induced by the ambient Euclidean metric and volume measure d*V*_*g*_. We observe points *x*_1_, …, *x*_*N*_ ∈ ℝ^*D*^ drawn i.i.d. from a distribution supported on (or in a tubular neighborhood of) ℳ . In the on-manifold case,

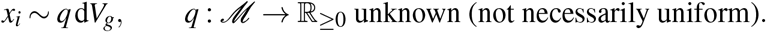

In the near-manifold case, *x*_*i*_ = *ξ*_*i*_ + *ε*_*i*_ with *ξ*_*i*_ ∼ *q* d*V*_*g*_ on ℳ and small ambient noise *ε*_*i*_ supported in a tubular neighborhood; when needed we write *π*(*x*_*i*_) ∈ ℳ for the nearest-point projection and assume *π*_#_*µ* = *q* d*V*_*g*_.

#### Overview

We adopt the manifold hypothesis in a form tailored to single-cell data: the observed profiles *x*_1_, …, *x*_*N*_ ∈ ℝ^*D*^ are sampled on (or very near) a *d*-dimensional Riemannian submanifold ℳ ⊂ ℝ^*D*^ according to an unknown, not-necessarily uniform sampling density *q* with respect to the manifold’s volume measure. Our goal is to recover informative, low-distortion coordinates that reflect the *intrinsic* geometry of ℳ rather than artifacts of the ambient space.

A principled way to expose this geometry is via the Laplace–Beltrami operator (LBO), Δ_*g*_, the natural generalization of the Laplacian to curved spaces. Its eigenfunctions act like a “Fourier basis” on ℳ : low-frequency eigenfunctions capture broad organization, while higher frequencies resolve finer structure. In practice, Δ_*g*_ is not observed directly; instead, methods such as Laplacian Eigenmaps and Diffusion Maps^9,10^ build a *graph Laplacian* from an affinity (similarity) graph on the samples. This graph operator converges to Δ_*g*_, especially when affinities are computed with *manifold-adaptive kernels* that correct for nonuniform sampling *q*^12^.

TopoMetry builds on this theory by constructing a *spectral scaffold*: an orthonormal eigenbasis of graph-LBO eigenvectors that provides data-driven, intrinsic coordinates. Each eigenvector can be read as a smooth “mode of variation” across cells, and the collection of the leading modes spans a coordinate system that encodes both local neighborhoods and global relationships. To avoid choosing a single resolution, TopoMetry forms a *multiscale spectral scaffold* by reweighting these modes across diffusion times (powers of the diffusion operator), thereby blending fine-grained and long-range structure in a single representation. The scaffold size is automatically selected from intrinsic-dimensionality estimates, so that subtle transcriptional signals and broad phenotypic organization are both retained without ad hoc parameter tuning.

Finally, TopoMetry reconstructs a neighborhood graph *in the scaffold space* and learns the corresponding graph LBO of the spectral scaffold itself—capturing the “geometry of the geometry.” The resulting operators (diffusion potentials and refined similarity graphs) serve as high-fidelity inputs for downstream analyses such as clustering, trajectory inference, pseudotime, and layout optimization (e.g. MAP, PaCMAP). In short, TopoMetry progressively denoises and sharpens manifold structure across scales, yielding coordinates that are stable, interpretable, and faithful to the intrinsic geometry of single-cell atlases.

#### Computational implementation

TopoMetry was designed for ease of use and computational efficiency. It integrates popular approximate nearest-neighbor (ANN) libraries such as NMSlib and hnswlib. Most graph operations are performed on sparse matrices and scale efficiently with multithreading. Each workflow step is implemented as a modular scikit-learn-style transformer, enabling custom pipelines. Core modules include estimators of neighborhood graphs, affinity kernels and operators, eigendecompositions, and layout optimizations. Analyses are orchestrated by the `TopOGraph` class. The modular design allows seamless integration with machine-learning and single-cell analysis pipelines. Additional utilities estimate intrinsic dimensionalities, evaluate embeddings, and visualize results. High-level APIs simplify usage: a single line of code can run a complete analysis and output a comprehensive report, including direct compatibility with `Scanpy` (^4^) and the broader Python ecosystem for single-cell analyses. The source code is available at https://github.com/davisidarta/topometry. Documentation and tutorials are available at https://topometry.readthedocs.io/en/latest/.

#### Assumed input

TopoMetry expects as input a cell-by-feature matrix that has been scaled or standardized, typically by Z-score normalization of gene expression. This preprocessing ensures that each feature contributes comparably to similarity calculations and avoids dominance by highly expressed or variable genes. A standardized input is also advantageous because it allows cell–cell similarity to be computed directly from feature correlations. In practice, this is implemented efficiently using cosine distances, which are equivalent to correlation distances on standardized data. In single-cell genomics, the input is usually subset to include only highly variable features, and we employed such standard preprocessing with default parameters in all demonstrated analyses.

#### Distance calculation

Distances are computed differently at each stage of the workflow. For building the initial spectral scaffold, we use cosine distance on normalized data, which is robust to differences in sequencing depth and scale and corresponds to the correlation distance when the input is standardized. The cosine (or correlation) distance between two cells *x*_*i*_, *x* _*j*_ ∈ ℝ^*d*^ is defined as:

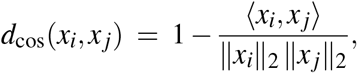

where ⟨·, ·⟩ denotes the Euclidean inner product.

For refined graphs built on the scaffold (*P*of_Z), we use Euclidean distance in the learned spectral coordinates, which provides a natural metric for diffusion geometry. The Euclidean distance between two spectral embeddings *z*_*i*_, *z* _*j*_ ∈ ℝ^*r*^ is defined as:

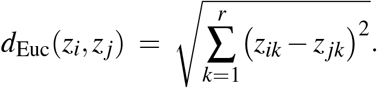

#### Kernels and affinity estimation

Let ℳ ⊂ ℝ^*M*^ be (near) a Riemannian submanifold with unknown sampling density *q*. For each sample *x*_*i*_ ∈ ℝ^*M*^, denote by nbrs(*x*_*i*_) the ordered set of its *k*-nearest neighbors under a chosen dissimilarity *d*(·, ·) (Euclidean or cosine-based; see “Cosine/correlation geometry” below). The affinity graph 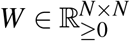 is built from locally rescaled distances and then symmetrized.

##### Adaptive scaling (per-point bandwidth)

For robustness to heterogeneous sampling, a local bandwidth *σ*_*i*_ is estimated from the median neighbor distance of *x*_*i*_:

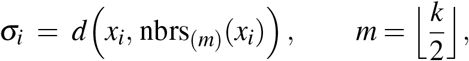

i.e. the *m*-th order statistic of {*d*(*x*_*i*_, *x* _*j*_) : *x* _*j*_ ∈ nbrs(*x*_*i*_)}. This *σ*_*i*_ encodes local crowding (small in dense regions, large in sparse ones) and stabilizes locality.

##### Cosine/correlation geometry and angularization

When using the cosine metric, we convert cosine *distance* 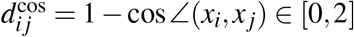 to an angle *θ*_*i j*_ ∈ [0, *π*],

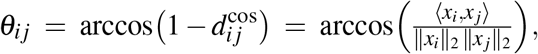

and use *d*_*i j*_ := *θ*_*i j*_ instead of the Euclidean distance. On feature-wise standardized (z-scored) inputs, cos∠(*x*_*i*_, *x* _*j*_) equals the Pearson correlation, so *θ*_*i j*_ = arccos(*ρ*_*i j*_) places similarities on the unit sphere (“correlation geometry”). Distances are numerically clipped to their natural ranges [0, *π*] (angular) or [0, 2] (cosine), and optionally squared (default), before kernelization.

##### Bandwidth-adaptive (variable) kernel

We first form a left-normalized, variable-bandwidth dissimilarity

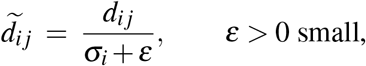

which rescales each row by its local scale *σ*_*i*_. The basic affinity is then

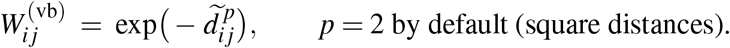

Finally, we symmetrize *W* ← (*W* + *W*^⊤^)/2. This row-wise scaling followed by symmetrization closely approximates classical (*σ*_*i*_, *σ*_*j*_)-variable bandwidth kernels while being simple and fast to compute.

##### Neighborhood density and “pseudomedian”

From 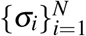 we derive a continuous density proxy *ω*_*i*_ by linearly mapping the interval [min _*j*_ *σ*_*j*_, max _*j*_ *σ*_*j*_] to [2, *k*]:

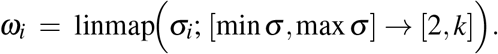

Large *ω*_*i*_ (near *k*) indicates sparsity; small *ω*_*i*_ (near 2) indicates locality within dense regions. This quantity can guide mild neighbor-set expansion in under-sampled areas (optionally replacing *k* with *k*^′^ ≥ *k*) and parameterizes the adaptive decaying kernel.

##### Adaptively decaying kernel (density-dependent tails)

To slow decay in sparse regions and sharpen it in dense ones, we make the decay exponent a function of *ω*_*i*_. Define

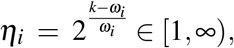

so that *η*_*i*_ ≈ 1 in sparse areas (*ω*_*i*_ ≈ *k*) and *η*_*i*_ grows as neighborhoods become denser.

Using the same left-normalized 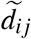, we set

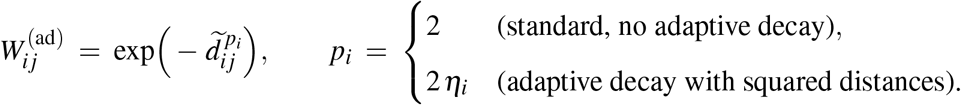

Equivalently, with squared distances disabled, one uses *p*_*i*_ = *η*_*i*_. The resulting kernel decays gently across sparse regions (preserving informative long-range neighbors) and more steeply inside dense regions (preventing oversmoothing). As above, we symmetrize *W* at the end.

##### Fixed-bandwidth (global) kernel

For comparison or when desired, a global scale *σ* > 0 yields

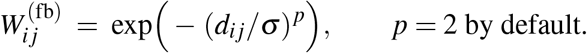

##### Summary

TopoMetry provides: (i) a fixed-bandwidth Gaussian/RBF kernel; (ii) a variable, per-point bandwidth kernel using median-*k* scaling; (iii) an optional neighborhood-expansion variant driven by the density proxy *ω*_*i*_; and (iv) an *adaptively decaying* kernel with density-dependent exponents. Distance preprocessing supports Euclidean geometry and correlation/cosine geometry via angularization. These design choices make affinity estimation sensitive to manifold structure while robust to heterogeneous sampling, providing a stable foundation for subsequent Laplace–Beltrami operator approximations and diffusion-based embeddings.

#### Intrinsic dimensionality estimation

Intrinsic dimensionality (i.d.) can be loosely defined as the minimum number of parameters needed to describe a high-dimensional system accurately. Multiple notions exist (e.g., local vs. global i.d.), and accurate estimation remains an active area of research. Estimating the global i.d. of a dataset is closely related to estimating the dimensionality of its underlying manifold, which is crucial when selecting the number of components for dimensionality reduction. Local i.d. is also useful as an auxiliary variable to characterize data geometry. Notably, the similarity kernels employed in TopoMetry are related to the Farahmand–Szepesvári–Audibert (FSA) estimator, leveraging ratios of distances to the *k*/2- and *k*-th nearest neighbors as a proxy for local sampling density.

##### Maximum Likelihood Estimator (MLE)

Consider i.i.d. observations *X*_*i*_ = *g*(*Y*_*i*_) that embed lower-dimensional samples *Y*_*i*_ drawn from an unknown density *f* via a continuous, smooth map *g*. For a point *x*, assume *f* (*x*) is approximately constant within a small sphere *S*_*x*_(*R*) of radius *R*, so observations in *S*_*x*_(*R*) form a homogeneous Poisson process. Replacing the radius by the *k*-nearest neighbors yields the local MLE for intrinsic dimension around *x*:

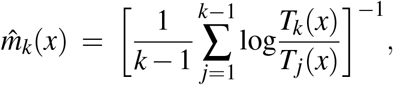

where *T*_*j*_(*x*) is the distance from *x* to its *j*-th nearest neighbor. A global estimate is obtained by averaging the local estimates over all *x*.

##### Manifold-adaptive dimensionality estimation (FSA)

The FSA method uses two nested neighborhoods around *x* (at *k* and *k*/2 neighbors) to estimate local i.d.:

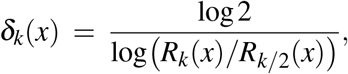

where *R*_*j*_(*x*) denotes the distance from *x* to its *j*-th nearest neighbor. The ratio *Rk*/*R*_*k*/2_ coincides with the scaling factor used in TopoMetry’s bandwidth-adaptive kernel. The global i.d. is reported as the median of the local *δk*(*x*) values.

#### Laplacian-type and diffusion operators

Let 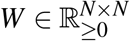 be a symmetric affinity (similarity) matrix built on the samples, with zero diagonal. Denote the (weighted) degree of node *i* by *d*_*i*_ = ∑ _*j*_ *W*_*ij*_ and *D* = diag(*d*1,…, *d*_*N*_ ). Graph Laplacians translate this similarity structure into linear operators whose spectra summarize geometry and connectivity; diffusion operators turn *W* into a Markov process on cells.

*Three standard graph Laplacians:*

(i) Unnormalized: *L* = *D* −*W*.
(ii) Symmetric normalized: *L*_sym_ = *D*^−1/2^*LD*^−1/2^ = *I* − *D*^−1/2^*WD*^−1/2^.
(iii) Random-walk (row-normalized): *L*_rw_ = *D*^−1^*L* = *I* − *D*^−1^*W*.

All three are positive semidefinite on a connected graph, with a trivial eigenpair (0, **1**). In practice, *L*_sym_ is convenient for symmetric eigensolvers and is less sensitive to degree heterogeneity; *L*_rw_ is directly linked to random walks and diffusion. Spectral embeddings such as Laplacian Eigenmaps^9^ use the eigenvectors associated with the smallest nonzero eigenvalues of one of these Laplacians (often dropping the constant eigenvector) as intrinsic coordinates.

##### Diffusion operator and diffusion time

Row-normalizing *W* yields the (row-stochastic) diffusion operator

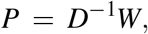

which defines a Markov chain on cells: (*P*^*t*^ )_*ij*_ is the probability of transitioning from cell *i* to cell *j* in *t* steps. The leading eigenvalues *λ*_1_ ≥ *λ*_2_ ≥ · · · ≥ 0 and right eigenvectors *ψ*_1_, *ψ*_2_, … of *P* provide *diffusion coordinates* at time *t*,

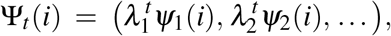

which emphasize multi-step connectivity and progressively suppress noise^10^. When a single *t* is not desired, one can use alternative scalings (e.g., “diffusion potential” mappings) that highlight long-range structure without fixing *t*.

##### Anisotropic (α) normalization to reduce sampling bias

Nonuniform sampling density can bias graph operators. A standard correction (Diffusion Maps) reweights *W* by powers of the degree before row-normalization:

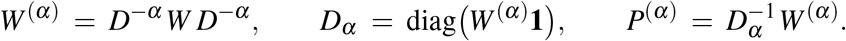

Here 0 ≤ *α* ≤ 1 tunes the level of debiasing: *α* = 0 leaves *W* unchanged; *α* = 1 largely removes effects of sampling density. A related “semi-anisotropic” variant applies the degree correction during normalization while retaining the original edge weights *W* ; this can stabilize spectra while preserving the original notion of similarity.

##### Symmetric diffusion operator for stable spectra

While *P*^(*α*)^ is generally asymmetric, it is similar to a symmetric matrix with the *same* eigenvalues:

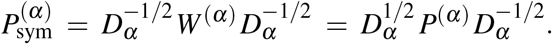

This Hermitian form is numerically well behaved; its eigenvectors can be mapped back to those of *P*^(*α*)^ by left-multiplication with 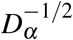. Diffusion-map embeddings are then obtained from the leading eigenpairs of *P*^(*α*)^ (or equivalently 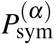), optionally raised to diffusion time *t*.

##### Connections to manifold geometry

Under standard assumptions (smooth manifold, appropriate kernels and bandwidths), graph Laplacians converge to the Laplace– Beltrami operator Δ_*g*_ on the data manifold, and *P*^*t*^ approximates heat diffusion 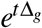^9,10^. This link justifies using Laplacian- and diffusion-based spectra as intrinsic, geometry-aware coordinates for downstream single-cell analysis.

#### Eigendecomposition, spectral scaffolds, and multiscaling

Let *P* ∈ ℝ^*N*×*N*^ be the diffusion operator derived from the affinity graph. We start from the eigendecomposition

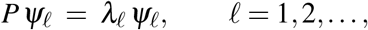

with eigenvalues ordered by magnitude 1 = *λ*_1_ ≥ *λ*_2_ ≥ · · · ≥ 0. For numerical stability it is often convenient to work with a symmetric similarity *P*_sym_ = *D*^1/2^*PD*^−1/2^, which is similar to *P* and therefore has the same spectrum; its eigenvectors are orthonormal in the Euclidean inner product and can be mapped back to right–eigenvectors of *P* by left multiplication with *D*^−1/2^.

##### Spectral scaffold

We define the *spectral scaffold* as

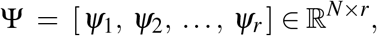

whose columns are mutually orthogonal (orthonormal after normalization) and ordered by decreasing |*λ*ℓ|. The leading column *ψ*_1_ is the trivial stationary mode (constant on each connected component) and is dropped in embeddings. The scaffold size *r* is chosen adaptively using intrinsic dimensionality estimates and eigengap heuristics, so it expands or contracts with dataset complexity rather than being fixed a priori.

##### Diffusion coordinates at timescale t

Geometry at a specific diffusion timescale *t* ∈ ℕ is obtained by reweighting scaffold columns by 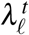 . For cell *i*,

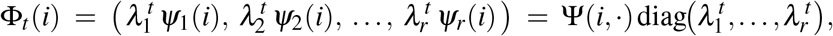

which progressively suppresses high-frequency (noise-like) components as *t* increases, emphasizing long-range organization.

##### Multiscale spectral scaffold

Besides using single *t*, TopoMetry builds a multiscale representation by analytically aggregating all diffusion times. Using the geometric-series identity 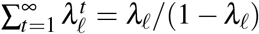 (valid for *λ*ℓ ∈ [0, 1) and after removing the trivial mode *λ*_1_ = 1), we form

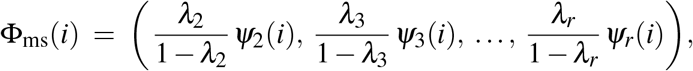

or, in matrix form,

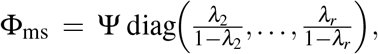

optionally restricting to eigenvalues *λ*ℓ > 0 to avoid numerical instabilities. This *multiscale spectral scaffold* is equivalent to concatenating diffusion coordinates across all times with exponentially decaying weights, thereby blending fine-grained neighborhoods (small *t*) and long-range connectivity (large *t*) in a single, compact set of coordinates.

##### Normalization and symmetry handling

When a symmetric similarity is used for eigensolving, the obtained eigenvectors are mapped to the right–eigenvectors of *P* via the similarity transform described above and then ℓ_2_-normalized, ensuring that columns of Ψ remain mutually orthogonal. With *r* selected adaptively, the resulting spectral scaffold (single–time Φ_*t*_ or multiscale Φ_ms_) provides geometry-aware coordinates that adjust to the intrinsic complexity of the single-cell atlas.

#### Refined graph

Given spectral coordinates *Z* ∈ ℝ^*N*×*r*^ (either a single-time Φ_*t*_ or the multiscale spectral scaffold Φ_ms_), we rebuild an affinity graph 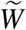 *in scaffold space* to sharpen geometry and suppress residual noise.

##### Construction in Z-space

We measure distances with the Euclidean metric *d*_*ij*_ =∥*z*_*i*_ − *z* _*j*_∥_2_. Using the same adaptive-bandwidth principle as above, each node *i* receives a local scale *σ*_*i*_ given by the distance to its median *k*-nearest neighbor in *Z*-space. Affinities are then computed by

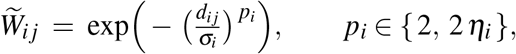

where *p*_*i*_ = 2 yields a bandwidth-adaptive Gaussian and *p*_*i*_ = 2 *η*_*i*_ is an adaptively decaying variant that slows decay in sparse regions and sharpens it in dense ones. The density proxy *ω*_*i*_ and decay factor 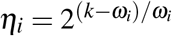 are defined as in the kernel section. Finally, we symmetrize

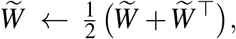

ensuring a real symmetric affinity suitable for spectral operators (with standard numerical guards).

##### Optional neighborhood expansion

In severely under-sampled areas, we allow a mild, density-aware increase of the neighbor set to *k*^′^ ≥ *k* (guided by *ω*_*i*_), recompute *σ*_*i*_on nbrs_*k*_′(*i*), and apply the same formulae.

From 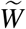 we compute the degree matrix 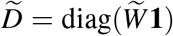 and derive Laplacian-type operators whose normalization yields refined diffusion operators. By default, TopoMetry applies anisotropic reweighting to these operators, producing a geometry-adapted transition matrix that serves as the backbone for downstream analyses. This refined diffusion operator is then used for tasks such as clustering, trajectory inference, pseudotime analysis, and visualization with graph layout algorithms, ensuring that all inferences are grounded in a refined representation of the original manifold of cell identities.

#### Layout optimization

For visualization, TopoMetry applies graph-layout optimization algorithms to the refined graphs. By default, we provide *TopoMAP* (an efficient UMAP-style layout applied to a precomputed manifold graph) and *TopoPaCMAP*; both are optimized on the refined diffusion operator and its neighborhood structure. Layout trajectories can be inspected to assess convergence and stability.

##### Uniform Manifold Approximation and Projection (UMAP) / Manifold Approximation and Projection (MAP)

UMAP-style objectives interpret a weighted graph as the 1-skeleton of a *fuzzy simplicial set* (FSS): an edge set with graded memberships *µ*_*ij*_ ∈ [0, 1] that encode how strongly *i* and *j* belong to one another’s neighborhood. Given a low-dimensional embedding *Y* = {*y*_*i*_}, a second fuzzy edge set with memberships *ν*_*ij*_(*Y* ) ∈ [0, 1] is induced by a fixed, monotonically decreasing function of the low-dimensional distance ∥*y*_*i*_ − *y* _*j*_∥. The layout is obtained by minimizing the cross-entropy between the two fuzzy edge sets:

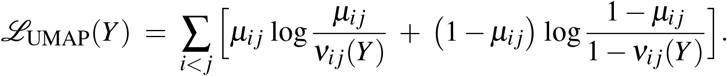

In TopoMetry, *MAP* denotes applying the same objective to an *arbitrary, precomputed* manifold graph (here, the refined graph in scaffold space); we refer to this implementation as *TopoMAP*.

##### Pairwise Controlled Manifold Approximation and Projection (PaCMAP)

PaCMAP optimizes a robust pairwise objective that balances three types of pair relationships: (i) *near pairs* (local neighbors), (ii) *mid-near pairs* (pairs connecting adjacent neighborhoods), and (iii) *further pairs* (distant non-neighbors). Let local scales be defined from the average distance to the 4th–6th nearest neighbors; write the high-dimensional, scale-adjusted distance as

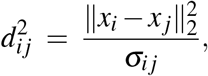

and let 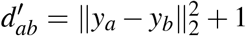 denote the corresponding low-dimensional distance surrogate. The PaCMAP loss is

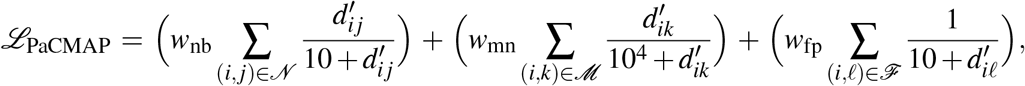

where 𝒩, ℳ, ℱ are the sets of near, mid-near, and further pairs selected per the original sampling rules, and (*w*nb, *w*_mn_, *w*fp) are stage-specific weights (e.g., Stage 1: (2, 1000, 1), Stage 2: (3, 3, 1), Stage 3: (1, 0, 1)). In TopoMetry, PaCMAP is initialized from the spectral scaffold and uses neighbor information from the refined manifold graph, yielding stable, high-fidelity layouts that respect both local neighborhoods and broader organization.

#### Riemann diagnostics and visualizations

Let *Y* ∈ ℝ^*N*×*m*^ be an embedding (typically *m* = 2 for visualization) with coordinate functions *y*^(*a*)^ : {1,…, *N*} → ℝ given by the *a*-th column of *Y*, and let *L* ∈ ℝ^*N*×*N*^ be a (symmetrized) graph Laplacian built on the refined affinity. We center the embedding, 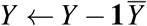, and write *Ly* for the discrete application of *L* to a vector *y*.

##### Discrete dual metric and embedding-space metric

For each sample *i* and coordinate indices *a, b* ∈ {1,…, *m*}, define the discrete *dual* metric (a covariant tensor) by

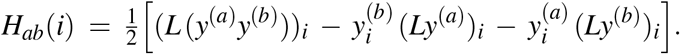

Stacking over *a, b* yields *H*(*i*) ∈ ℝ^*m*×*m*^. We obtain the embedding-space Riemannian metric *G*(*i*) as the (regularized) pseudoinverse of *H*(*i*):

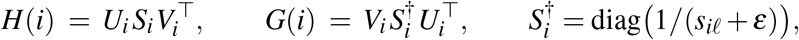

with a small *ε* > 0 to stabilize near-zero singular values. Finally, we project *G*(*i*) to the symmetric positive definite cone by eigenvalue clipping and normalize its overall scale (e.g. unit trace) to make ellipse sizes comparable across points.

##### Local distortion indicatrix

Let 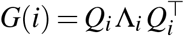 with Λ_*i*_ = diag(*λi*,1 ≥ · · · ≥ *λ*_*i,m*_ > 0). The *indicatrix* at *i* is the ellipse (for *m* = 2)

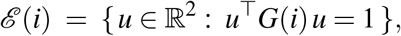

whose principal axes have directions given by the columns of *Q*_*i*_ and semi-axis lengths

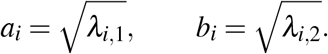

Thus 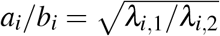 quantifies local anisotropy (directional stretching), while det *G*(*i*) = *λi*,1*λi*,2 quantifies local area change. In plots, we scale (*a*_*i*_, *b*_*i*_) by a global factor and ensure each ellipse fits within the axes (conservative circumscribed-circle bound) to avoid clipping.

##### Contraction/expansion scalar field

We color points (or ellipses) by a centered log-determinant of the metric,

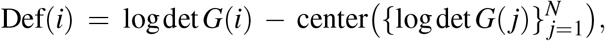

with the center chosen as the median (robust) or mean, so that Def > 0 indicates local *expansion* and Def < 0 indicates *contraction*. Optionally, we smooth this scalar over the graph by *t* steps of diffusion, Def ← *P*^*t*^ Def, where *P* = *D*^−1^*W* is the random-walk operator derived from *L* = *D* −*W* ; we then re-center and use robust percentile clipping with symmetric limits for color normalization.

##### Visualization modes

(i) *Localized indicatrices:* draw ellipses ℰ (*i*) for a uniform subset of points, optionally modulating their size by a normalized anisotropy score log(*λi*,1/*λi*,2). The underlying scatter may be colored by Def(*i*) or other annotations. (ii) *Grid-averaged field:* build a regular grid over the embedding, and at each grid site *g* average nearby metrics, 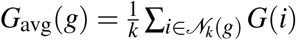 draw one ellipse from *G*_avg_(*g*) and color it by the grid-averaged Def. A thinning step enforces a minimum separation between grid sites to limit overlap.

##### Geometric interpretation

If *φ* : ℳ → ℝ^*m*^ denotes the embedding map, then *G*(*i*) estimates the pullback metric 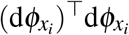 (up to the discrete Laplacian normalization), so that logdet 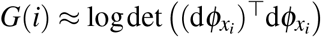 reports local area change and the eigenstructure of *G*(*i*) reports directional distortion. The plotted indicatrices therefore act as *distortion rulers*, revealing where a 2-D layout expands, contracts, or shears the intrinsic geometry.

#### Data imputation

We perform diffusion-based imputation on the cell–cell graph. Let *X* ∈ ℝ^*N*×*G*^ denote the cell-by-gene matrix to be imputed, and let *P* ∈ ℝ^*N*×*N*^ be the Markov (row-stochastic) diffusion operator built on the refined manifold graph. For a diffusion time *t* ∈ ℕ, the imputed matrix is the graph diffusion

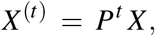

i.e. each gene is smoothed across the graph while preserving its marginal scale.

##### Automatic selection of the diffusion time

To avoid oversmoothing, we select *t* from a candidate grid 𝒯 = {*t*_1_, …, *t*_*m*_} by contrasting the gene–gene correlation structure of *X*^(*t*)^ against a null in which cross-cell associations have been destroyed but univariate marginals are preserved. Concretely, we first identify a small panel of highly variable genes to score. Let *X*_top_ ∈ ℝ^*N*×*g*^ be the submatrix of these genes. For each *t* ∈ 𝒯 :

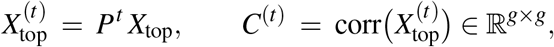

and we define a scalar score as the mean absolute off-diagonal correlation,

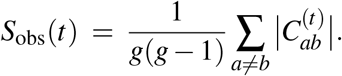

##### Null model and empirical test

For each *t*, we generate a null distribution by independently permuting the rows of every column of *X*_top_ (gene-wise shuffling across cells), diffusing with the same *t*, and recomputing the score:

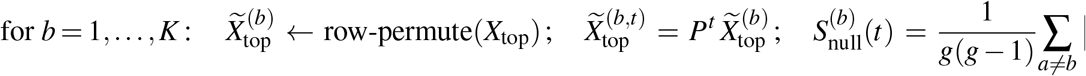

We then compute an empirical *p*-value and a *z*-score:

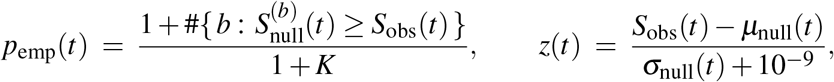

with *µ*_null_, *σ*_null_ the mean and standard deviation of 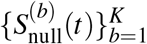 . The selected time

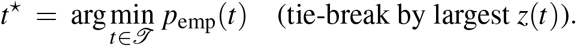

After identifying an optimal diffusion time t, the optimal imputation result is:

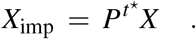

This procedure selects the weakest amount of smoothing that yields gene–gene correlation structure significantly exceeding what diffusion would induce under a null with broken cross-cell associations. In doing so, it explicitly guards against the common pitfall of oversmoothing, where excessive diffusion can erase genuine biological heterogeneity and introduce spurious correlations that mimic structure but are in fact artifacts. By benchmarking the observed correlation profiles against an empirical null, the method ensures that only structure reproducibly supported by the data is retained. This design minimizes the risk of false positives: correlations that arise purely from graph connectivity rather than underlying transcriptional programs are filtered out, since they also appear in the null distribution and therefore fail the statistical test. At the same time, genuine patterns of co-expression that reflect biological coordination between genes are amplified, as they persist in the observed data but not under randomized permutations. The approach thus balances denoising with fidelity to the original data, producing imputations that are statistically principled, biologically meaningful, and less likely to bias downstream analyses such as clustering, trajectory inference, or differential expression.

#### Signal filtering

TopoMetry provides a graph-based low-pass filter to denoise sample-level signals while respecting manifold structure. Let *s* ∈ ℝ^*N*^ be a per-cell signal and *P* ∈ ℝ^*N*×*N*^ the row-stochastic diffusion operator on the refined graph (default: built in the multiscale spectral scaffold space). The filtered signal after *t* diffusion steps is

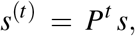

which corresponds to heat-kernel smoothing on the graph: increasing *t* attenuates high-frequency (noisy, rapidly varying) components and preserves low-frequency structure aligned with the topology.

##### Signal construction

Users supply a column in the sample annotations. If the column is categorical, we form a binary indicator

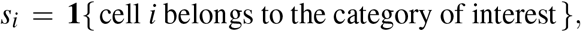

else a numeric column is used directly (non-finite entries set to 0). For stress tests, one may optionally add pre-filter Gaussian noise, *s* ← *s* + *σε* with *ε* ∼ 𝒩 (0, *I*) and *σ* ≥ 0.

##### Choice of operator and time

Filtering can use the Markov operator derived in any space (`msZ`, `Z`, `X`); the default `msZ` leverages the refined multiscale geometry. The parameter *t* ∈ ℕ controls smoothness: *t* = 0 returns the original signal, small *t* mildly denoises within neighborhoods, and large *t* approaches the stationary distribution (oversmoothing). In practice, modest values (e.g. *t* ≈ 4−8) preserve biological contrasts while reducing noise.

##### Normalization (optional)

After filtering, an optional rescaling can be applied. If the signal is intended as a probability or indicator, one can keep the native range. Otherwise, a min–max normalization maps *s*^(*t*)^ to [0, 1]:

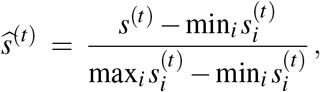

guarded for degenerate ranges. This facilitates downstream visualization and thresholding.

For reproducibility, both the raw vector and its filtered counterpart are recorded in the sample metadata, including the diffusion time *t* and the operator choice. The procedure is computationally light (matrix–vector products, compatible with sparse *P*) and can be applied to categorical labels, risk scores, or any continuous per-cell quantity to obtain geometry-aware smoothed estimates.

### 4.3 Quantification of geometry preservation

To systematically evaluate how well different representations preserve the intrinsic geometry of single-cell data, we developed a set of four complementary metrics. Each metric operates directly on the diffusion operators associated with the original data space and the candidate embedding, thereby enabling a scale-aware and operator-native assessment of geometry preservation.

#### Row-neighborhood F1 score (PF1)

Let *P*_*x*_ and *P*_*y*_ denote two row-stochastic diffusion operators to be compared. For each cell *i*, let *N*_*x*_(*i*) and *N*_*y*_(*i*) denote the sets of the top-*k* neighbors defined by the largest transition probabilities in *P*_*x*_ and *P*_*y*_, respectively.

The per-row F1 score is given by

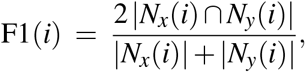

and the PF1 score is obtained as the average over all cells:

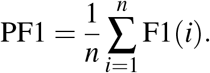

This score captures the preservation of local neighborhood identities, independently of edge weights.

#### Row-wise Jensen–Shannon similarity (PJS)

To compare the transition probability distributions themselves, we use the Jensen–Shannon (JS) divergence. For each row *i*, let *p*_*i*_ and *q*_*i*_ denote the probability vectors corresponding to the nonzero transitions of *P*_*x*_ and *P*_*y*_. The JS divergence between *p*_*i*_ and *q*_*i*_ is

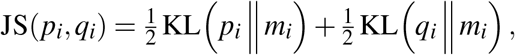

where 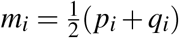 and KL denotes the Kullback–Leibler divergence. We then define the similarity as

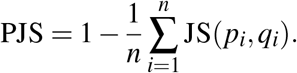

Unlike PF1, this metric is sensitive to the weights of transitions, thus evaluating the fidelity of local transition probabilities.

#### Spectral Procrustes alignment (SP)

Diffusion operators admit an eigendecomposition *Pψ*_ℓ_ = *λ*_ℓ_*ψ*_ℓ_, where {*λ*_ℓ_, *ψ*_ℓ_} are eigenpairs ordered by |*λ*_ℓ_|. For a given timescale *t*, the diffusion map is defined as

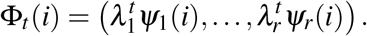

Given the diffusion coordinates 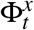 and 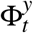 from *P*_*x*_ and *P*_*y*_, we align them via orthogonal Procrustes:

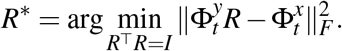

The alignment quality is quantified by the coefficient of determination,

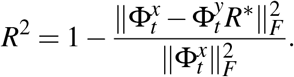

The SP score is averaged across multiple diffusion times *t*, capturing agreement of the meso- and global-scale geometry of the manifold.

#### Rationale for metric selection

These three metrics together capture complementary aspects of geometry preservation across scales: PF1 and PJS quantify fidelity of local neighborhoods, and SP evaluates alignment of diffusion eigencoordinates at mesoscopic scales. Compared to previously used metrics, they provide a more faithful evaluation of single-cell representations. Global scores based on PCA depend strongly on graph-layout initialization rather than underlying geometry. Geodesic correlation, though sensitive to structure, overweights large-scale distances and becomes biased in the presence of multiple disjoint submanifolds, as typical in single-cell data. Clustering- or label-based scores are unreliable in real-world settings where ground-truth identities are unknown. Finally, Riemannian diagnostics such as distortion heatmaps and indicatrices are restricted to 2-D embeddings, whereas single-cell analyses require latent spaces with tens or hundreds of dimensions. Our operator-native metrics therefore offer a principled, scalable, and interpretable quantification of geometry preservation across learned representations.

### 4.4 Standard analysis workflow

Single-cell analysis workflows have converged on a widely adopted pipeline that couples dimensionality reduction by Principal Component Analysis (PCA) with graph construction and visualization using Uniform Manifold Approximation and Projection (UMAP). This approach underpins the majority of published studies and is implemented as the default in most software ecosystems for single-cell genomics.

#### Preprocessing

Raw count matrices are typically subjected to two standard preprocessing steps. First, counts are normalized for library size (total molecules per cell) and log-transformed to stabilize variance. Second, a subset of highly variable genes (HVGs) is selected, usually between 1,000 and 5,000, under the assumption that these capture the majority of biologically relevant variation while reducing technical noise. The resulting HVG matrix is then standardized (Z-score normalized) so that each gene has mean zero and unit variance, preventing highly expressed genes from dominating subsequent analyses.

#### Principal Component Analysis

The standardized HVG matrix is projected into a lower-dimensional space using PCA. The number of retained components is typically chosen heuristically (e.g. 30–100) based on an *ad hoc* “elbow point”, rather than based on intrinsic dimensionality. The Euclidean distances between cells in this PCA space are then used to build a *k*-nearest-neighbors (kNN) graph, which serves as the foundation for clustering and visualization.

#### UMAP graph layout

The kNN graph is converted into an affinity graph using fuzzy simplicial sets, which UMAP optimizes into a two-dimensional embedding. The resulting map aims to preserve local neighborhoods while maintaining a global structure that is visually interpretable. This 2-D representation is widely used for cluster annotation, lineage inference, and exploratory analysis, despite being optimized for visualization rather than strict geometric fidelity.

#### clustering and identification of marker genes

Clustering was performed with the Leiden community detection algorithm using default parameters via `scanpy.tl.leiden`. To ensure fair comparisons, we applied the same clustering algorithm with identical `resolution` parameters across all evaluated workflows, including TopoMetry. Marker genes were identified with `scanpy.tl.rank_genes_groups` after constructing dendrograms on the representation associated with the clustering results, using the logistic regression method (`method=‘logreg’`), which has been shown to achieve optimal results in single-cell data^39^.

#### Summary

Together, these steps—log and size normalization, HVG selection, Z-score standardization, PCA projection, kNN graph construction, and UMAP embedding—constitute the de facto standard in single-cell analysis. While computationally efficient and easy to implement, the workflow is based on strong assumptions: that global variance identifies biologically meaningful structure, that a fixed number of PCs captures the latent geometry, and that UMAP embeddings accurately reflect manifold relationships. These assumptions have rarely been tested systematically, motivating the development of alternative frameworks such as TOPOMETRY.

#### RNA velocity analysis

RNA velocity analysis was performed using the `scVelo` toolkit^40^, with all steps executed using default parameters. The exception was the calculation of neighborhood graphs and moments, which by default rely on the PCA latent space, but were instead computed using the TopoMetry multiscale spectral scaffold. For TopoMetry analysis, spliced and unspliced counts were aggregated, and default preprocessing was applied to provide a standardized matrix of gene expression as input.

#### T cell receptor analysis

The ECCITE-TCR dataset was obtained with precomputed TCR analysis and metadata. For the TICA dataset, T cell receptor (TCR) clonotype analysis was performed using the `scirpy` toolkit for immune receptor analysis in Python^33^. TCR similarities were computed using the alignment of amino acid sequences. Default parameters were used for all remaining steps.

#### Variational inference

Variational autoencoder analyses were performed using the `scvi-tools` framework^20,21^. Models were trained on GPU using default parameters, including preprocessing steps handled internally by the package. Latent spaces were computed and used to construct neighborhood graphs for Leiden clustering and UMAP visualizations.

### 4.5 Public datasets

We assembled a corpus of publicly available single-cell RNA-seq collections from the CellxGene Census (release `2023-07-25`). For each target `collection_id`, we queried the Census metadata table (`census_info/datasets`) to obtain the corresponding `dataset_id`s and then retrieved primary RNA measurements (`is_primary_data = True`) as AnnData objects separately for *Homo sapiens* and *Mus musculus*. Observational annotations included tissue-level fields (e.g., `tissue`, `tissue_general`, `assay`, `cell_type`). Individual datasets within a collection were concatenated (outer join) after loading. To ensure basic quality, we removed cells with fewer than 100 detected genes and genes expressed in fewer than 5 cells prior to downstream analysis. For computational tractability, collections with more than 10^5^ cells were excluded from this benchmarking pass, as were a small number of collections exhibiting upstream data issues (e.g., malformed or inconsistent matrices). For each processed collection we stored a compact result bundle containing: collection identifiers and names, cell and gene counts, a set of two-dimensional embeddings, and a results blob with evaluation tables and diagnostics. Additional datasets analysed in this manuscript are also publicly available: The murine pancreas development dataset was obtained through the `scVelo` toolkit for RNA velocity estimation and visualization. The PBMC68k dataset was freely downloaded from 10X Genomics. Additional PBMC datasets were obtained from the accession codes phs002048.v1.p1 (LES), GSE154386 (Dengue) and GSE138266 (MS). The paired RNA and TCR datasets were made publicly available by their authors and were obtained from Zenodo and from the Human Cell Atlas.

### 4.6 Performance benchmark

All methods were applied to the same preprocessed matrices (highly variable genes, library-size normalization, log(1+*x*), scaling), as described in the preceding section. We evaluated geometry-preservation performance across representations using a unified routine that computes neighborhood- and diffusion-based scores.

Concretely, for each collection:

- We computed a standard PCA model (up to 200 components or the maximal feasible given *n*_cells_ and *n*_genes_) and retained:
  1. the PCA space (`X_PCA`),
  2. UMAP on PCA neighborhoods (`X_PCA_UMAP`),
  3. UMAP computed directly on the scaled expression space with cosine distance (`X_UMAP_on_X`).
- We trained an `SCVI` model on raw counts (default `scvi-tools` settings), extracted the latent representation (`X_scVI`), and computed UMAP on its neighborhood graph (`X_UMAP_on_scVI`).
- We ran TopoMetry via `tp.sc.fit_adata`, which builds a first kernel on expression, performs an eigendecomposition to obtain a spectral scaffold, and refines the kernel on that scaffold before producing low-dimensional projections (e.g., TopoMAP/TopoPaCMAP). The fitted `TopOGraph` exposes Markov operators *P*_*X*_, *P*_*Z*_, and *P*_ms*Z*_ used in evaluation.

We assessed geometry preservation for *all* representations present in `adata.obsm` using the same parameters: Euclidean distance, local neighborhood size *k*=30, HNSW-based approximate nearest-neighbors search, and diffusion times *t* ∈ {1, 4, 8} with spectral rank *r*=64. The evaluation routine reports a table of metrics, including row-neighborhood *k*-NN overlap (PF1), row-wise Jensen–Shannon similarity (PJS), and Spectral Procrustes alignment (SP). PCA diagnostics recorded per collection included the full variance-explained spectrum and its cumulative sum. Default hyperparameters were used for all evaluated workflows to enable a fair performance comparison across datasets.

## Data and Code Availability

`topometry` is freely accessible as a Python library under the MIT license. The source code is available at https://github.com/davisidarta/topometry, and can be installed through the Python Package Index (PyPI) at https://pypi.org/project/topometry/. The library is extensively documented at https://topometry.readthedocs.io. Any additional information can be requested from the authors and will be made available upon request.

## Acknowledgments

We thank Leland McInness, Dmitry Kobak, and Akshay Agrawal for valuable comments on the ideas presented in this manuscript. We also thank Bruno Loyola Barbosa for assistance in testing early implementations of `topometry`.

