## Supplementary figures for "TopoMetry systematically learns and evaluates the latent geometry of single-cell data"

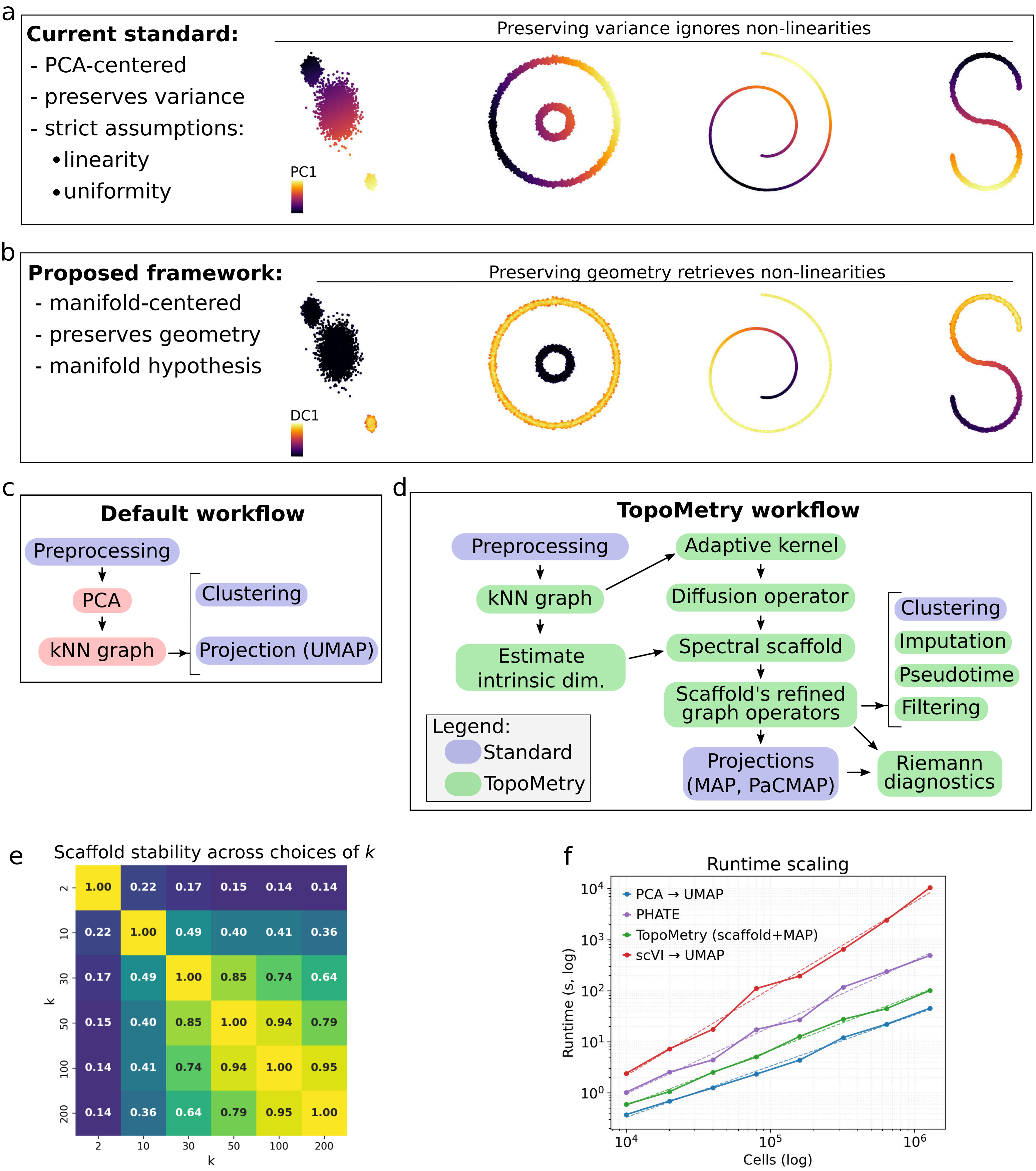
a
Current standard:
- PCA-centered
- preserves variance
- strict assumptions:
•linearity
•uniformity
Preserving variance ignores non-linearities
PC1
b
Proposed framework:
- manifold-centered
- preserves geometry
- manifold hypothesis
Preserving geometry retrieves non-linearities
DC1
c
Default workflow
Preprocessing
PCA
kNN graph
Clustering
Projection (UMAP)
d
TopoMetry workflow
Preprocessing
kNN graph
Estimate intrinsic dim.
Adaptive kernel
Diffusion operator
Spectral scaffold
Scaffold's refined graph operators
Projections (MAP, PaCMAP)
Riemann diagnostics
Clustering
Imputation
Pseudotime
Filtering
Legend:
Standard
TopoMetry
e
Scaffold stability across choices of k
k: 2, 10, 30, 50, 100, 200
2: 1.00 0.22 0.17 0.15 0.14 0.14
10: 0.22 1.00 0.49 0.40 0.41 0.36
30: 0.17 0.49 1.00 0.85 0.74 0.64
50: 0.15 0.40 0.85 1.00 0.94 0.79
100: 0.14 0.41 0.74 0.94 1.00 0.95
200: 0.14 0.36 0.64 0.79 0.95 1.00
k
f
Runtime scaling
PCA → UMAP
PHATE
TopoMetry (scaffold+MAP)
scVI → UMAP
Runtime (s, log)
Cells (log)
$10^0$ $10^1$ $10^2$ $10^3$ $10^4$
$10^4$ $10^5$ $10^6$

**Figure S1: Schematic overview of the current standard and the proposed framework for single-cell analysis.**

(a) Left: the current standard is centered around PCA, focused on preserving variance, and implicitly carries strict assumptions on the underlying geometry and sample distribution. Right: two-dimensional synthetic datasets colored by their first principal component (PC). Note how PCA fails to capture non-linear relationships and local structure. (b) Left: the proposed framework is centered on manifold learning, focuses on preserving geometry, and assumes only the manifold hypothesis. Right: two-dimensional synthetic datasets colored by their first diffusion component (DC). Note how spectral analysis successfully retrieves the underlying geometry regardless of non-linearity or uneven sampling. (c) Schematic of the current default workflow. After preprocessing, the PCA representation is used to learn a k-NN graph, which is then utilized for downstream analyses, such as clustering and visualization/projection. (d) Schematic of the TopoMetry workflow. After preprocessing, a kNN graph is learned from the original high-dimensional space and used to estimate intrinsic dimensionalities (I.D.) and cell–cell similarities (through an adaptive kernel), which are used to learn Laplacian-type and diffusion operators from the data. These operators are decomposed into spectral scaffolds, and a second, refined iteration of graph operators is built on the scaffold itself. These refined graphs are faithful representations of the manifold of cellular identities and can be utilized for downstream tasks, including clustering, visualization, imputation, pseudotime estimation, signal filtering, and manifold diagnostics. (e) Correlation matrix showing the agreement between scaffolds learned with different numbers of k-nearest neighbors across choices of $k$. Note how results remain stable for reasonable ranges of $k$ (30–100). (f) Runtime benchmark of TopoMetry against the current PCA$\rightarrow$UMAP standard, PHATE, and scVI across different dataset sizes. While less efficient than the PCA$\rightarrow$UMAP workflow, TopoMetry is computationally more scalable than alternative approaches.

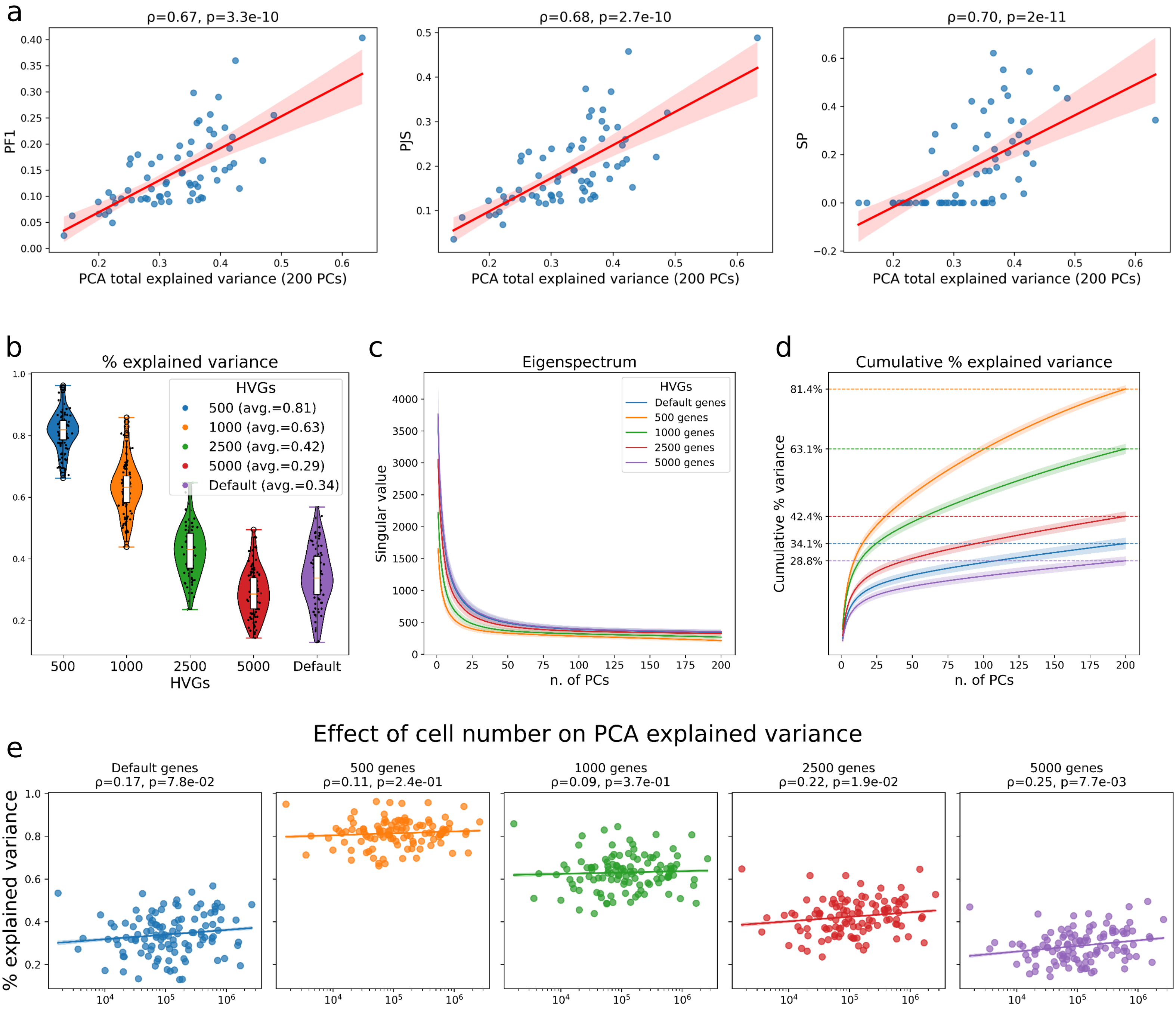
a
ρ=0.67, p=3.3e-10
PF1
PCA total explained variance (200 PCs)
ρ=0.68, p=2.7e-10
PJS
PCA total explained variance (200 PCs)
ρ=0.70, p=2e-11
SP
PCA total explained variance (200 PCs)
b
% explained variance
HVGs
500 (avg.=0.81)
1000 (avg.=0.63)
2500 (avg.=0.42)
5000 (avg.=0.29)
Default (avg.=0.34)
500
1000
2500
5000
Default
HVGs
c
Eigenspectrum
HVGs
Default genes
500 genes
1000 genes
2500 genes
5000 genes
Singular value
n. of PCs
d
Cumulative % explained variance
81.4%
63.1%
42.4%
34.1%
28.8%
Cumulative % variance
n. of PCs
e
Effect of cell number on PCA explained variance
Default genes
ρ=0.17, p=7.8e-02
500 genes
ρ=0.11, p=2.4e-01
1000 genes
ρ=0.09, p=3.7e-01
2500 genes
ρ=0.22, p=1.9e-02
5000 genes
ρ=0.25, p=7.7e-03
% explained variance

**Figure S2: Systematic Evaluation of PCA-Explained Variance Across Single-Cell Datasets.** (a) PCA scores in geometry-preservation metrics correlate with its total explained variance across datasets. (b) PCA total explained variance across scRNA-seq datasets depends on the selection of highly variable genes, decreases with larger numbers of genes, and is considerably low at default settings. (c) "Scree" plot of singular values, showing a stable "elbow point" at 30–50 PCs (higher with larger HVGs). (d) Cumulative percentage of explained variance across datasets and number of PCs, highlighting that the poor performance of PCA in scRNA-seq data cannot be attributed to an insufficient number of PCs. (e) PCA performance improves as the number of cells increases, particularly when considering larger numbers of HVGs.

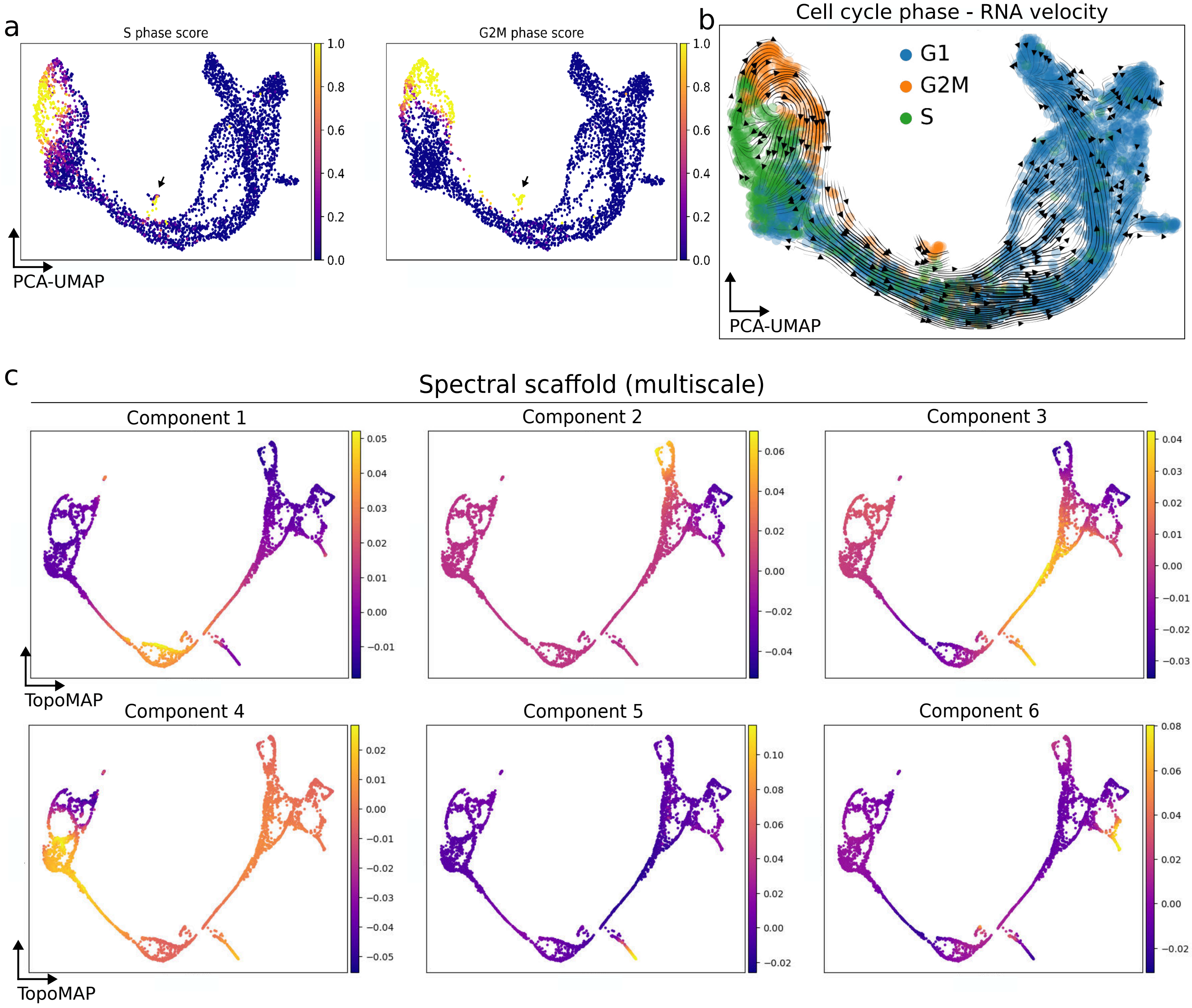
a
S phase score
G2M phase score
PCA-UMAP
b
Cell cycle phase - RNA velocity
G1
G2M
S
PCA-UMAP
c
Spectral scaffold (multiscale)
Component 1
Component 2
Component 3
Component 4
Component 5
Component 6
TopoMAP
TopoMAP

**Figure S3: The current PCA→UMAP standard fails to represent cell cycle geometry.** (a–b) PCA→UMAP visualizations of the murine pancreas development dataset, colored by inferred scores of different phases of the cell cycle (a), and the predicted cell cycle phase for each cell with RNA velocity overlay (b). The scores, predictions, and RNA velocity are highly suggestive of underlying cell cycle geometry, which is ignored and distorted by the PCA→UMAP visualization. (c) TopoMAP visualizations of the same dataset, colored by the first six components of the spectral scaffold. Note how each component encodes a different aspect of the underlying geometry.

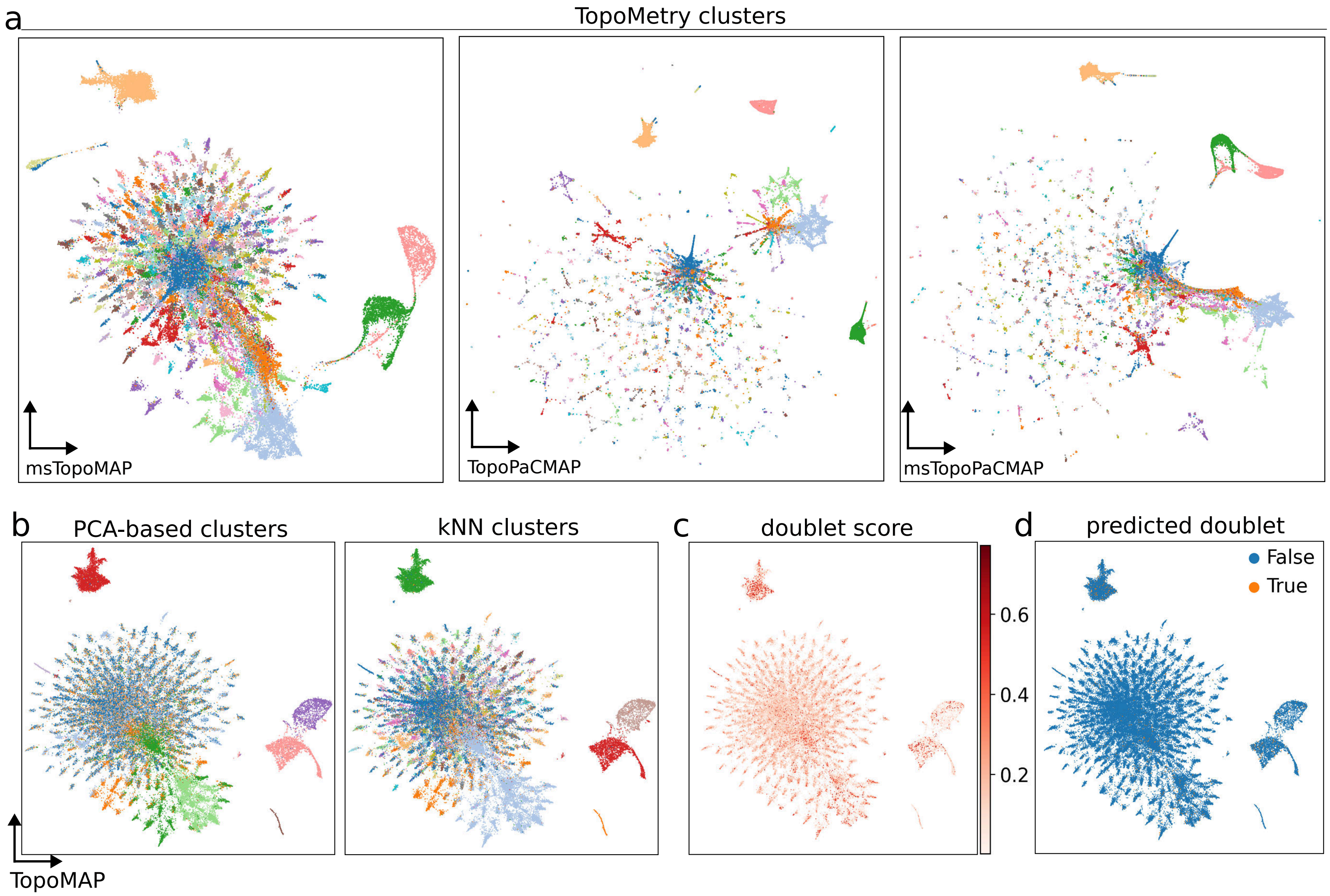
a
TopoMetry clusters
msTopoMAP
TopoPaCMAP
msTopoPaCMAP
b
PCA-based clusters
kNN clusters
TopoMAP
c
doublet score
0.6
0.4
0.2
d
predicted doublet
False
True

**Figure S4: TopoMetry's visualizations consistently detect T cell diversity.**
(a) Additional TopoMetry visualizations of the pbmc68k dataset (from left to right: MAP on the multiscale scaffold's operator, PaCMAP on the fixed-time or multiscale scaffold's operator). (b) TopoMAP visualizations colored by clustering results obtained from the standard PCA-based workflow (left) or with a standalone kNN graph (right). Note how the clustering results from the standalone kNN graph partly agree with TopoMetry's results and succeed in detecting some of the T cell clusters identified by TopoMetry. (c–d) TopoMAP visualizations colored by Scrublet's doublet score (c) and by predicted doublets (d), showing that the additional T cell clusters identified by TopoMetry cannot be attributed to doublets.

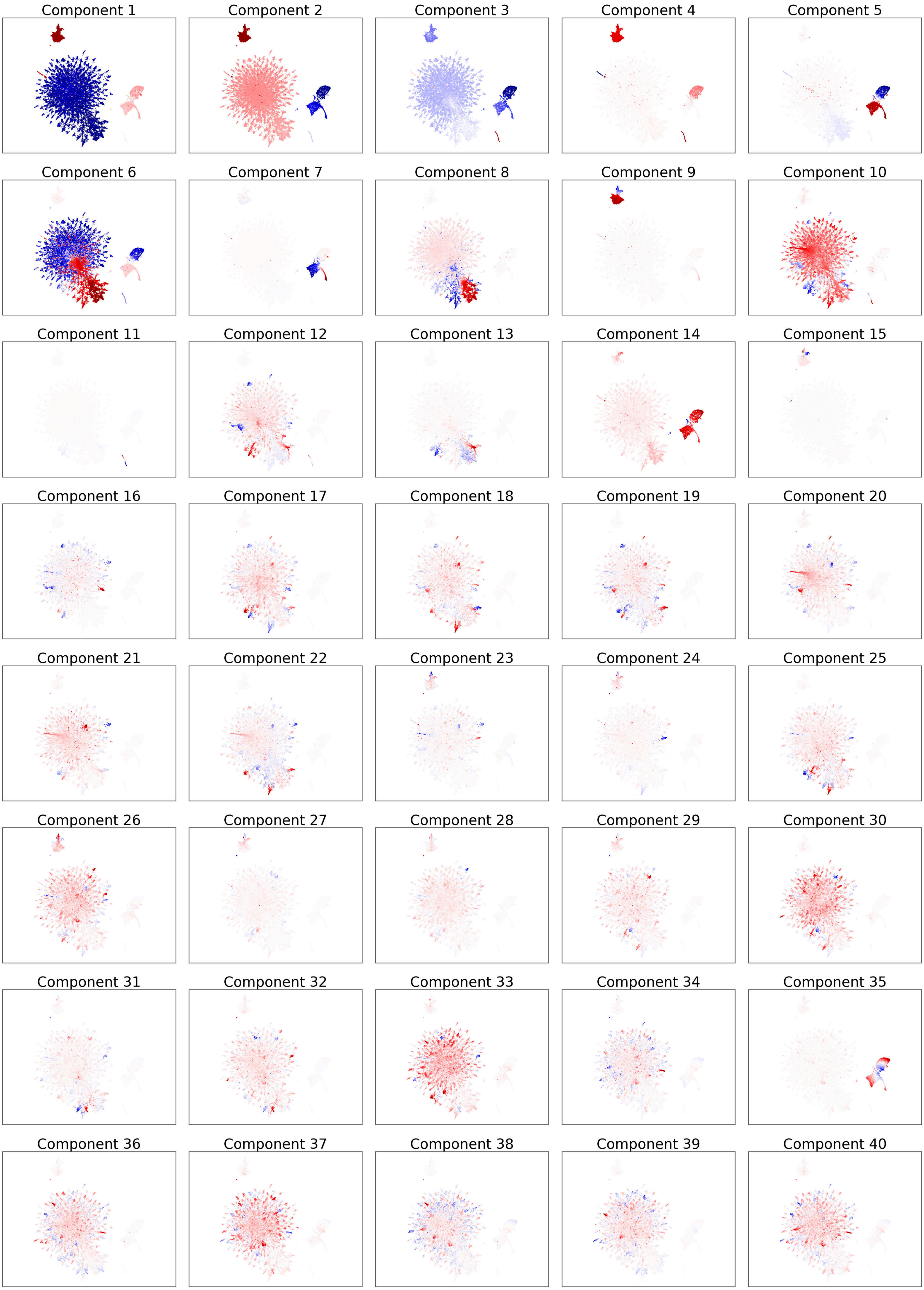
Component 1
Component 2
Component 3
Component 4
Component 5
Component 6
Component 7
Component 8
Component 9
Component 10
Component 11
Component 12
Component 13
Component 14
Component 15
Component 16
Component 17
Component 18
Component 19
Component 20
Component 21
Component 22
Component 23
Component 24
Component 25
Component 26
Component 27
Component 28
Component 29
Component 30
Component 31
Component 32
Component 33
Component 34
Component 35
Component 36
Component 37
Component 38
Component 39
Component 40

**Figure S5: The spectral scaffold of PBMCs.**
Panel of the first 40 components of the spectral scaffold learned from the pbmc68k dataset by TopoMetry. Note how the information encoded by the first few components is associated with global class separation and long-range relationships. The encoded information becomes increasingly detailed as the number of components increases. TopoMetry estimates intrinsic dimensionalities and uses an ad hoc eigengap to automatically identify the ideal number of scaffold components to be constructed.

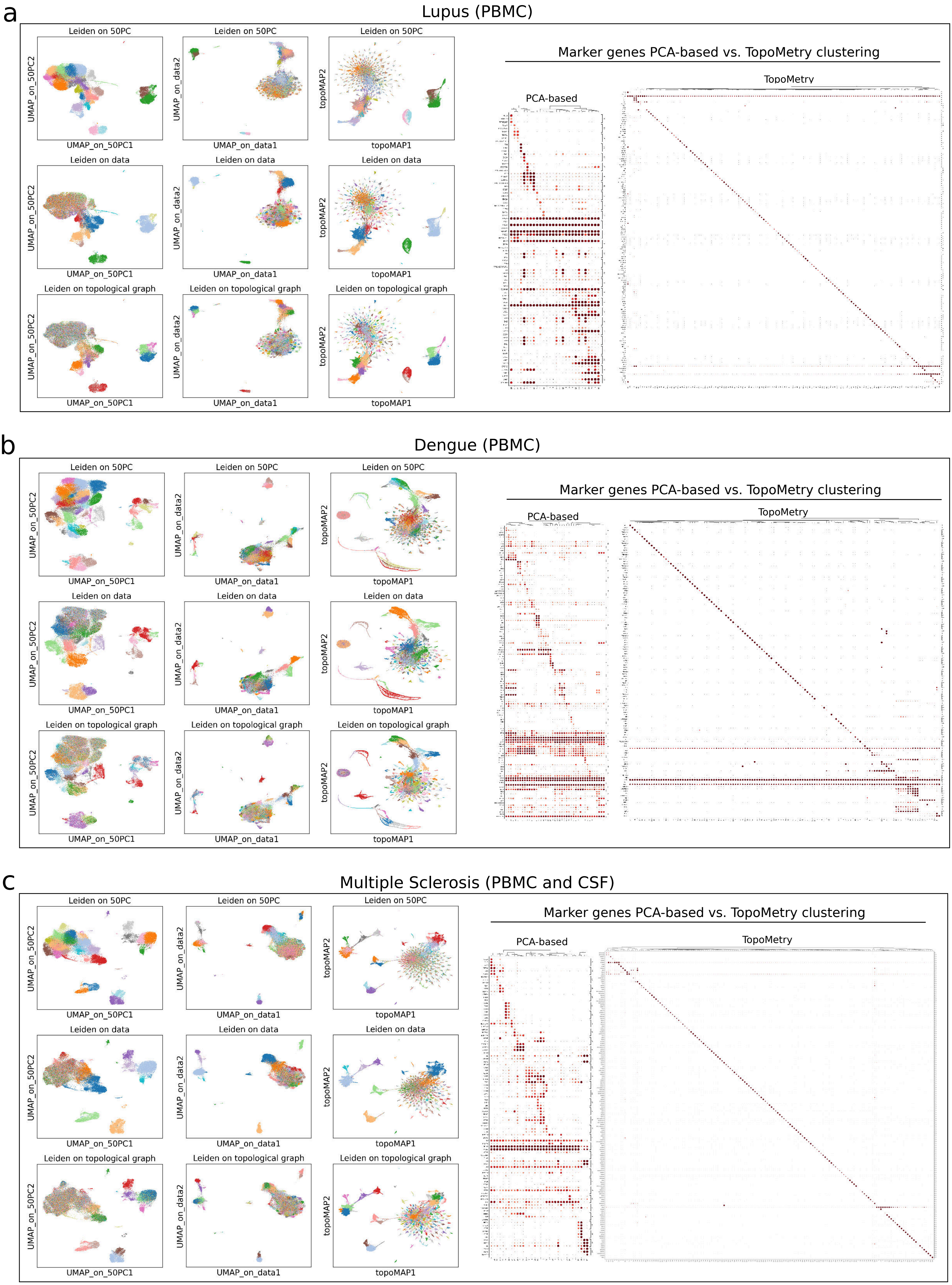

a
Lupus (PBMC)
Leiden on 50PC
Leiden on data
Leiden on topological graph
UMAP_on_50PC1
UMAP_on_50PC2
UMAP_on_data1
UMAP_on_data2
topoMAP1
topoMAP2
Marker genes PCA-based vs. TopoMetry clustering
PCA-based
TopoMetry
b
Dengue (PBMC)
Leiden on 50PC
Leiden on data
Leiden on topological graph
UMAP_on_50PC1
UMAP_on_50PC2
UMAP_on_data1
UMAP_on_data2
topoMAP1
topoMAP2
Marker genes PCA-based vs. TopoMetry clustering
PCA-based
TopoMetry
c
Multiple Sclerosis (PBMC and CSF)
Leiden on 50PC
Leiden on data
Leiden on topological graph
UMAP_on_50PC1
UMAP_on_50PC2
UMAP_on_data1
UMAP_on_data2
topoMAP1
topoMAP2
Marker genes PCA-based vs. TopoMetry clustering
PCA-based
TopoMetry

**Figure S6: Transcriptional diversity of T cells across diseases.**
A set of three panels, each containing visualization and clustering results for a dataset of peripheral mononuclear blood cells (PBMC) using (i) the PCA→UMAP workflow (50 PCs retained), (ii) standalone kNN graph and UMAP ("on data"), and (iii) TopoMetry, in addition to the marker gene dotplots for the clusters found by (i) and (iii). Across all datasets, the PCA-based results suggest little T cell diversity and present non-specific marker gene signatures, standalone results identify some additional T cell populations, and TopoMetry results detect the full range of T cell clusters with highly specific marker genes. (a) Panel for the Lupus dataset, comprising PBMCs from healthy donors and systemic lupus erythematosus patients. (b) Panel for the Dengue dataset, consisting of PBMCs collected from a dengue fever patient. (c) Panel for the multiple sclerosis dataset, comprising PBMCs and mononuclear cells from cerebrospinal fluid (CSF) samples from healthy donors and multiple sclerosis patients.

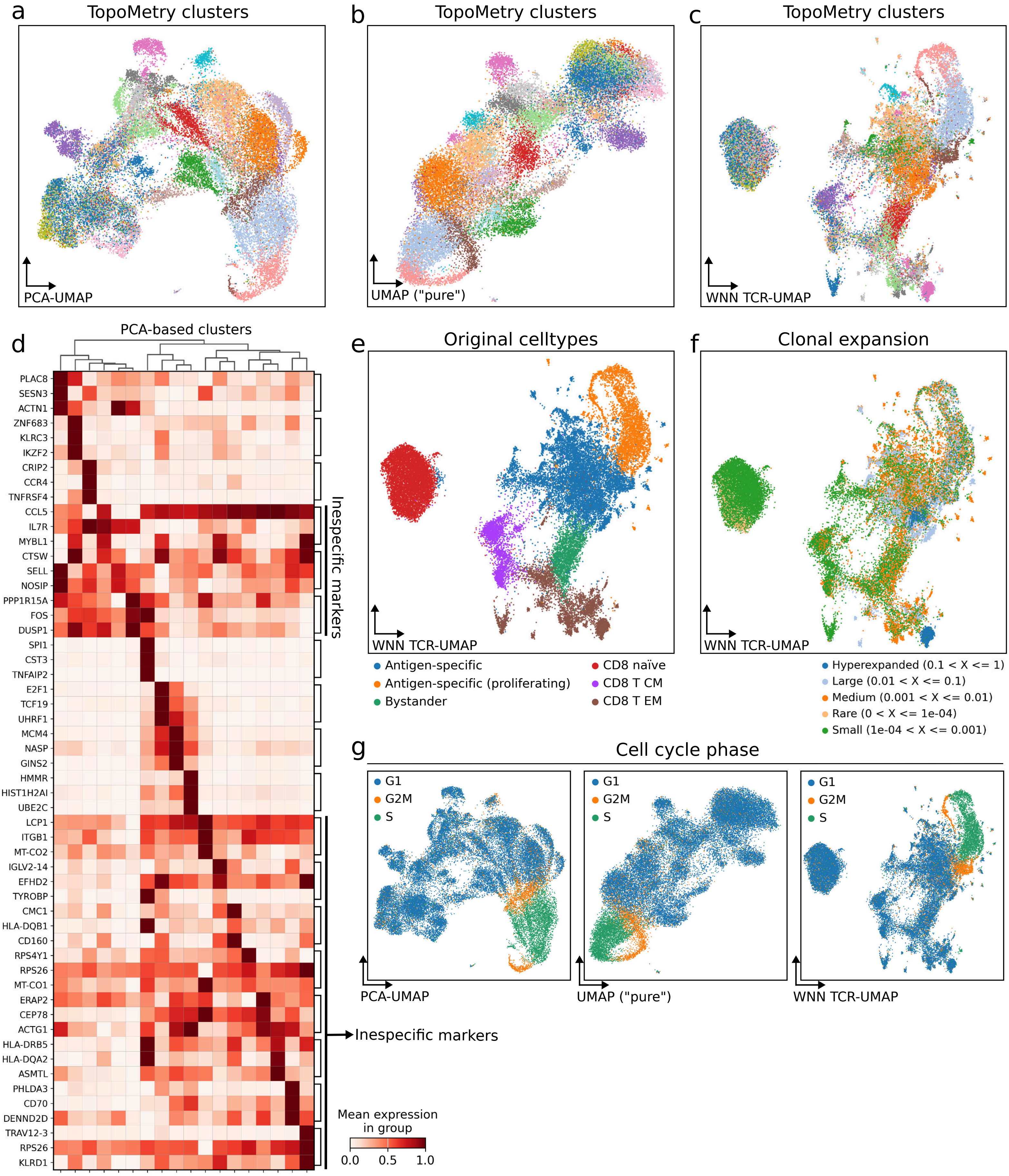
a
TopoMetry clusters
PCA-UMAP
b
TopoMetry clusters
UMAP ("pure")
c
TopoMetry clusters
WNN TCR-UMAP
d
PCA-based clusters
PLAC8
SESN3
ACTN1
ZNF683
KLRC3
IKZF2
CRIP2
CCR4
TNFRSF4
CCL5
IL7R
MYBL1
CTSW
SELL
NOSIP
PPP1R15A
FOS
DUSP1
SPI1
CST3
TNFAIP2
E2F1
TCF19
UHRF1
MCM4
NASP
GINS2
HMMR
HIST1H2AI
UBE2C
LCP1
ITGB1
MT-CO2
IGLV2-14
EFHD2
TYROBP
CMC1
HLA-DQB1
CD160
RPS4Y1
RPS26
MT-CO1
ERAP2
CEP78
ACTG1
HLA-DRB5
HLA-DQA2
ASMTL
PHLDA3
CD70
DENND2D
TRAV12-3
RPS26
KLRD1
Inespecific markers
Inespecific markers
Mean expression in group
0.0
0.5
1.0
e
Original celltypes
WNN TCR-UMAP
Antigen-specific
Antigen-specific (proliferating)
Bystander
CD8 naïve
CD8 T CM
CD8 T EM
f
Clonal expansion
WNN TCR-UMAP
Hyperexpanded (0.1 < X <= 1)
Large (0.01 < X <= 0.1)
Medium (0.001 < X <= 0.01)
Rare (0 < X <= 1e-04)
Small (1e-04 < X <= 0.001)
g
Cell cycle phase
G1
G2M
S
PCA-UMAP
UMAP ("pure")
WNN TCR-UMAP

**Figure S7: Additional comparisons on the ECCITE-TCR dataset.**
2-D visualizations of the ECCITE-TCR dataset using (a) PCA→UMAP, (b) standalone UMAP ("pure UMAP"), and (c) a weighted nearest-neighbors (WNN) graph built using both RNA and TCR information, all colored by TopoMetry's clustering results. Note how the paired TCR–RNA approach succeeds in detecting some of the additional clusters of $CD8^{+}$ TEM and TCM cells identified by TopoMetry. (d) Matrixplot of the top 3 marker genes found for PCA-based clustering results, with non-specific markers highlighted. (e–f) Paired WNN visualizations colored by original cell type annotations (e) and clonal expansion (f). (g) Same visualizations as in (a–c), colored by predicted cell cycle phase. Note how all visualizations fail to successfully represent the cell cycle geometry.

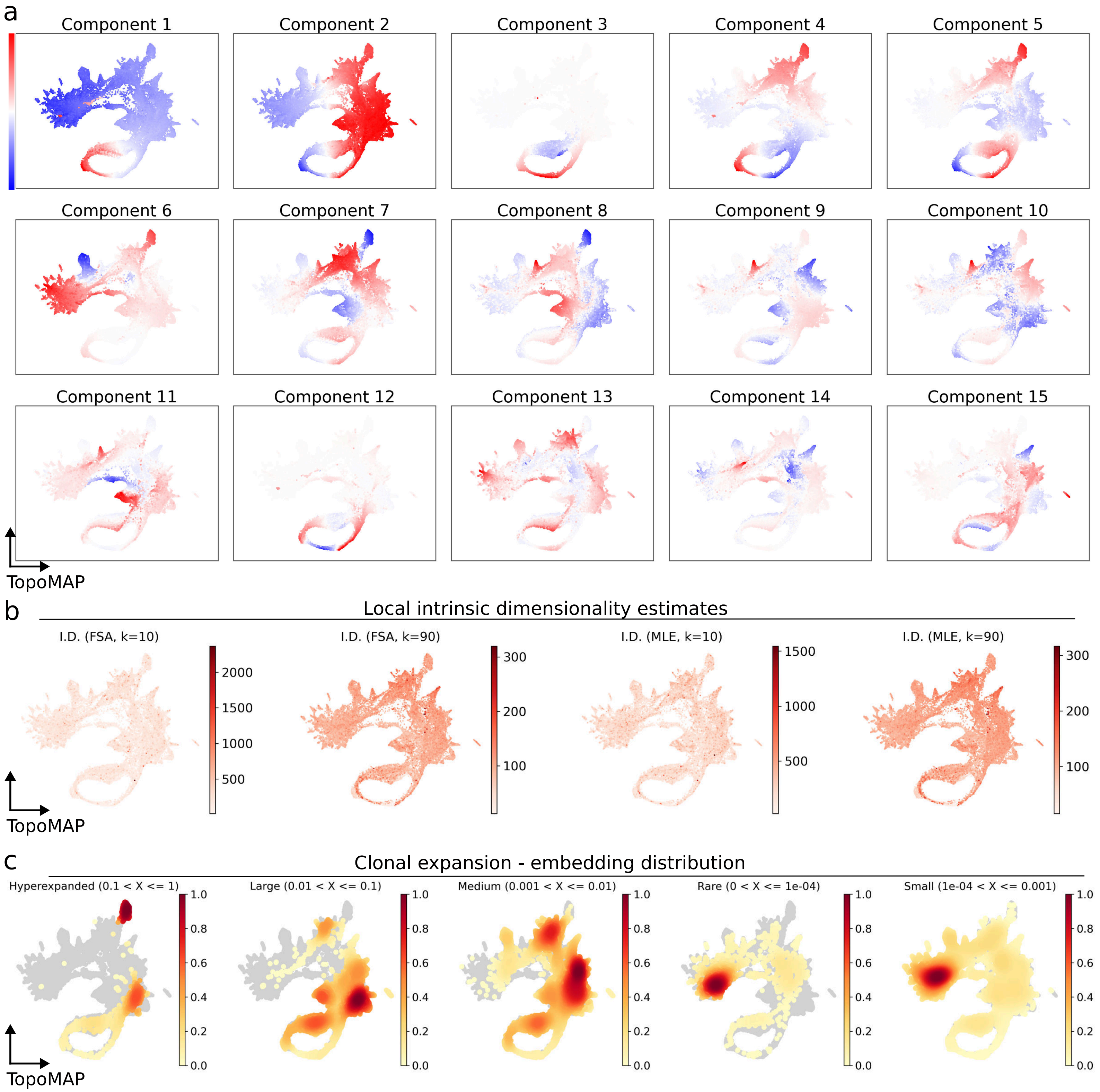
a
Component 1
Component 2
Component 3
Component 4
Component 5
Component 6
Component 7
Component 8
Component 9
Component 10
Component 11
Component 12
Component 13
Component 14
Component 15
TopoMAP
b
Local intrinsic dimensionality estimates
I.D. (FSA, k=10)
I.D. (FSA, k=90)
I.D. (MLE, k=10)
I.D. (MLE, k=90)
TopoMAP
c
Clonal expansion - embedding distribution
Hyperexpanded (0.1 < X <= 1)
Large (0.01 < X <= 0.1)
Medium (0.001 < X <= 0.01)
Rare (0 < X <= 1e-04)
Small (1e-04 < X <= 0.001)
TopoMAP

**Figure S8: Geometrical properties of clonal expansion dynamics.**
(a) Panel of TopoMAP visualizations of the ECCITE-TCR dataset, colored by the first 15 components of the spectral scaffold. Note how the first few components describe the global geometry (e.g., the lineage inference from proliferating cells to mature TEM and TCM and antigen-specific hyperexpanded T cells), while the following components progressively add local information to the scaffold. (b) TopoMAP visualizations, colored by estimated local intrinsic dimensionality (I.D.) using different methods (FSA, MLE) and choices of $k$-neighbors, with homogeneous estimates across different regions of the manifold. (c) TopoMAP visualizations, colored by the distribution density of clone sizes across the embedding.

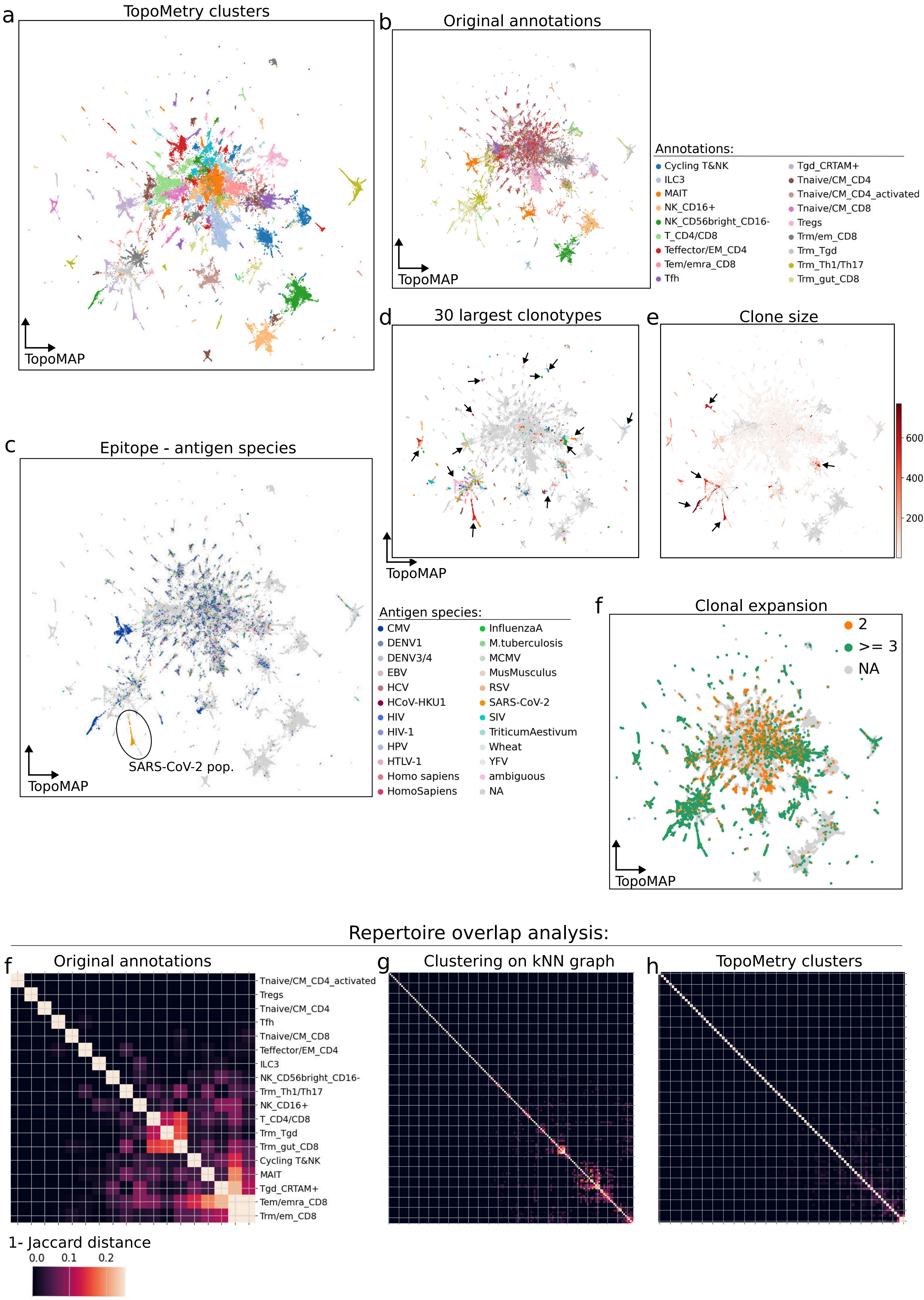
a
TopoMetry clusters
TopoMAP
b
Original annotations
TopoMAP
Annotations:
Cycling T&NK
ILC3
MAIT
NK_CD16+
NK_CD56bright_CD16-
T_CD4/CD8
Teffector/EM_CD4
Tem/emra_CD8
Tfh
Tgd_CRTAM+
Tnaive/CM_CD4
Tnaive/CM_CD4_activated
Tnaive/CM_CD8
Tregs
Trm/em_CD8
Trm_Tgd
Trm_Th1/Th17
Trm_gut_CD8
c
Epitope - antigen species
SARS-CoV-2 pop.
TopoMAP
d
30 largest clonotypes
TopoMAP
e
Clone size
600
400
200
Antigen species:
CMV
DENV1
DENV3/4
EBV
HCV
HCoV-HKU1
HIV
HIV-1
HPV
HTLV-1
Homo sapiens
HomoSapiens
InfluenzaA
M.tuberculosis
MCMV
MusMusculus
RSV
SARS-CoV-2
SIV
TriticumAestivum
Wheat
YFV
ambiguous
NA
f
Clonal expansion
2
>= 3
NA
TopoMAP
Repertoire overlap analysis:
f
Original annotations
Tnaive/CM_CD4_activated
Tregs
Tnaive/CM_CD4
Tfh
Tnaive/CM_CD8
Teffector/EM_CD4
ILC3
NK_CD56bright_CD16-
Trm_Th1/Th17
NK_CD16+
T_CD4/CD8
Trm_Tgd
Trm_gut_CD8
Cycling T&NK
MAIT
Tgd_CRTAM+
Tem/emra_CD8
Trm/em_CD8
g
Clustering on kNN graph
h
TopoMetry clusters
1- Jaccard distance
0.0
0.1
0.2

**Figure S9: T cell clusters identified exclusively by TopoMetry are associated with specific clonotypes.**

Analysis of the T cell compartment of the Tissue Immune Cell Atlas (TICA) dataset, for which both RNA-seq and VDJ-seq data are available. (a–f) TopoMAP visualization of TICA's T cell compartment, colored by (a) TopoMetry's clusters; (b) original cell type annotations; (c) origin species of epitopes recognized by each cell's TCR based on amino acid sequence, with a TopoMetry cluster corresponding to a clonotype cluster that specifically recognizes SARS-CoV-2 antigens; (d) the largest 30 clonotypes detected by TCR amino acid sequence, highlighting that these clonotypes correspond to TopoMetry's encoded geometry; (e) clone size, highlighting the largest clones and their agreement with the proposed geometry-aware representations; and (f) clonal expansion, showing TopoMetry's clusters that correspond to hyperexpanded clones. (g–i) Repertoire overlap analysis quantifying the overlap between TCR sequences between different clusters across clustering results, considering (g) the original cell type annotations, (h) the results from the standalone kNN graph, and (i) TopoMetry's clustering results. Note how the overlap can be mitigated by avoiding the use of PCA-based clusters (as in g) and using a standalone graph approach instead (as in h), and how it is almost entirely abolished by the geometry-aware analysis (as in i).
